# Polymerization force-regulated actin filament-Arp2/3 complex interaction dominates self-adaptive cell migrations

**DOI:** 10.1101/2023.04.15.536869

**Authors:** Xindong Chen, Yuhui Li, Ming Guo, Bowen Xu, Yanhui Ma, Hanxing Zhu, Xi-Qiao Feng

## Abstract

Cells migrate by adapting their leading-edge behaviours to heterogeneous extracellular microenvironments (ECMs) during cancer invasions and immune responses. Yet it remains poorly understood how such complicated dynamic behaviours emerge from millisecond-scale assembling activities of protein molecules, which are hard to probe experimentally. To address this gap, we established a spatiotemporal “resistance-adaptive propulsion” theory based on the protein interactions between Arp2/3 complexes and polymerizing actin filaments, and a multiscale dynamic modelling system spanning from molecular proteins to the cell. Combining spatiotemporal simulations with experiments, we quantitatively find that cells can accurately self-adapt propulsive forces to overcome heterogeneous ECMs via a resistance-triggered positive feedback mechanism, dominated by polymerization-induced actin filament bending and the bending-regulated actin-Arp2/3 binding. However, for high resistance regions, resistance triggered a negative feedback, hindering branched filament assembly, which adapts cellular morphologies to circumnavigate the obstacles. Strikingly, the synergy of the two opposite feedbacks not only empowers cells with both powerful and flexible migratory capabilities to deal with complex ECMs, but also endows cells to use their intracellular proteins efficiently. In addition, we identify that the nature of cell migration velocity depending on ECM history stems from the inherent temporal hysteresis of cytoskeleton remodelling. We also quantitatively show that directional cell migration is dictated by the competition between the local stiffness of ECMs and the local polymerizing rate of actin network caused by chemotactic cues. Our results reveal that it is the polymerization force-regulated actin filament-Arp2/3 complex binding interaction that dominates self-adaptive cell migrations in complex ECMs, and we provide a predictive theory and a spatiotemporal multiscale modelling system at the protein level.

## Introduction

Cells migrate through coupling their propulsive forces generated by assembling cytoskeletons to extracellular microenvironments (ECMs) (1–4). Actin-based lamellipodia protrusion is a powerful force-generating system that drives cell migrations during the cancer invasion, immune surveillance, and embryonic development (1, 5–9). Arp2/3 complex binds on an existing actin filament and then nucleates a daughter filament, assembling into branched actin networks in lamellipodia and invadopodia (10, 11). The polymerization of the network generates a pushing force to open a sufficient wide channel in ECM to drive individual or collective cell migrations (12–14).

Clinical studies show that Arp2/3 complex-medicated migration is tightly associated with cancer invasion (15–20), and patients over expressing Arp2/3 complex have poor survivals in lung (16), breast(17), pancreatic(20), and colorectal cancers(15). In addition, the migrations of immune cells, such as dendritic cells and T cells, in three-dimensional ECMs are also extensively driven by the Arp2/3 complex formed lamellipodial protrusions (6, 7). However, ECMs *in vivo* are highly mechanically heterogeneous(21, 22). Both invasive cancer cells and immune cells need to migrate long distances to establish new tumors(21, 23) and to find killing targets(7, 24), respectively. Experimental studies show that the magnitude of ECM resistance can affect the density of lamellipodial branched actin filaments(1, 5, 11, 25), and the lamellipodial leading-edge velocity exhibits resistance-history dependent(1, 26). In addition, cells predominantly migrate along the path of least resistance in heterogeneous extracellular microenvironments (HTECMs)(6). All these studies suggested that the leading edge of migratory cells actively mechano-senses variations of ECMs and adapts its migrating behaviour (Fig. 1a)(1, 5, 6, 27) in response to complex ECMs. However, the migratory leading edge involves highly dynamic interplays of various proteins, including Arp2/3 complexes, actin monomers, actin filaments, Wiskott-Aldrich syndrome proteins (WASPs), ATP, capping proteins, leading-edge membrane, integrin-based adhesions and ECMs(5, 10, 11, 28). Due to both the temporal and spatial cross-scale complexities(29) (Fig. 1a), live imaging protein behaviours at millisecond and nanometre scales is intrinsically difficult(30). Thus, quantitatively interpreting how these complicated cell-scale self-adaptive migration behaviours (Fig. 1a) emerge from the dynamic activities of molecular proteins has been a grand and long-lasting challenge (31, 32). It not only seriously hampers our in-depth mechanistic understandings of *in vivo* cell migrations, but also hinders us from discovering target proteins for designing new medicines and gene editing therapies to prevent cancer cells from invasions or enhance immune cell infiltrations for solid cancer immunotherapy.

**Fig. 1.**
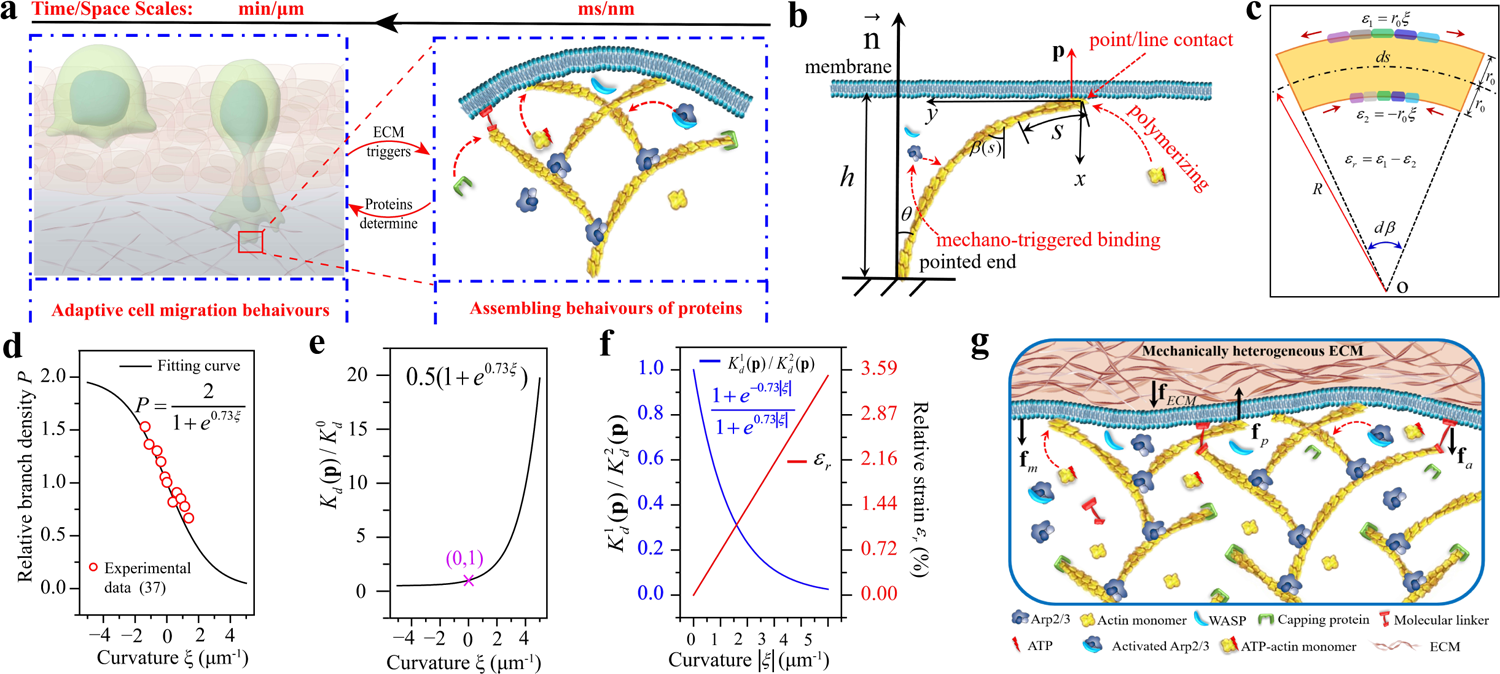
Polymerizing branched actin filaments at lamellipodial leading edge drive cells to migrate in ECMs. (**a**) Cell migrations in ECMs are spatial and temporal cross-scale biophysical behaviours performed by proteins. (**b**) Demonstration of the two-dimensional mechanical interaction for theoretical analysis. (**c**) Polymerization-induced deformation analysis of a small segment *ds* of actin filament. The convex side surface is stretched with a strain *ε*_1_ = *r*_0_*ξ* while the concave side surface is compressed with a strain *ε*_2_ = −*r*_0_*ξ* where *r*_0_ is the radius of actin filaments. The colour bar regions denote the binding surfaces of Arp2/3 complex on the convex and concave sides of the actin filaments. (**d**) Relationship between relative branched density and actin filament curvature from experimental data (37). It is fitted by an inverted sigmoid function, which is more reasonable than a linear fitting because the relative branched density should be neither minus nor excessively high. The relative branched density *P* is the ratio of the number of branch points with different curvatures to the corresponding number of mother actin filaments. (**e**) Relationship between the relative dissociation constants *K_d_* (**p**) *K_d_*^0^ and actin filament curvature where *K_d_* (**p**) and *K*^0^ are the dissociation constants in the bending and straight states, respectively (Supplementary information). (**f**) The relative dissociation constants *K_d_*^1^ (**p**) *K_d_*^2^(**p**) shows that the binding affinity of Arp2/3 complex on the convex side (*K*^1^ (**p**)) of a bending actin filament is much higher than the concave surface (*K_d_* ^2^ (**p**)). (**g**) Forces acting on the leading-edge membrane when a cell is migrating in ECMs. **f** *_p_* is the propulsive force generated by the polymerization of a branched actin filament. **f***_ECM_* is the extracellular resistance from the ECM. **f***_m_* is the tension force of the top and bottom lamellipodial membrane. **f***_a_* is the attachment force of a molecular linker, which links actin filament and the leading-edge membrane.

Constructing a predictive spatiotemporal multiscale modelling system that describes cellular dynamic behaviours from the molecular to cellular scales at the intersection of biology, physics, chemistry and computer science will greatly accelerate biomedical advancements(33, 34). From the level of molecular protein-protein interactions, we here derive a spatiotemporal ‘resistance-adaptive propulsion’ (RAP) theory based on the geometric nonlinear deformation mechanisms of polymerizing actin filaments and the mechano-chemical assembling behaviours of Arp2/3 complexes. The theory describes both spatial and temporal mechanical interactions between the polymerizing branched actin filaments and the bent leading-edge membrane constrained by ECM. On the basis of this RAP theory, we develop a multiscale spatiotemporal modelling system, which encompasses dynamic actin polymerization, capping protein inhabiting actin filament polymerization, nonlinear deformation of actin filaments, actin monomer diffusion, ATP binding on actin monomers and Arp2/3 complex, mechano-chemical assembly of Arp2/3 complex, detachments of molecular linkers, bent leading-edge membrane and HTECMs. It can not only simulate dynamic cell migrations in complex environments but also quantitatively shed light on synchronous interacting and assembling behaviours of multiple proteins. Combining spatiotemporal simulations with experiments tracking actin dynamics during cell migration, we quantitatively discovered a resistance-triggered positive feedback mechanism at the protein level for adapting propulsive forces to overcome the heterogeneity in ECM, and a resistance-triggered negative feedback mechanism for adapting cell morphology to circumnavigate regions with high ECM resistance. Strikingly, the combination of the two opposite feedback mechanisms along the broad lamellipodial leading edge shows formidable synergistic effects, empowering cells with both powerful and flexible migratory capabilities to deal with complex ECMs, and meanwhile endowing intracellular ATP resource with an optimal efficiency in fuelling cell migration. These insights explain why it is hard to prevent metastasis by cancer cells once they have acquired invasive ability. By monitoring the assembling behaviours of proteins, we further find that the nature of migration velocity depending on the resistance history of ECM is derived from the temporal hysteresis of adaptive actin cytoskeleton remodelling. In addition, we reveal that directional cell migration is dictated by the competition between the local stiffness of ECMs and the local polymerizing rates of the leading-edge actin cytoskeleton in response to gradients of chemotactic cues. Overall, we establish the first, to our knowledge, spatiotemporal biophysical theory and multiscale modelling system, which can accurately predict self-adaptive cell migrations in complex ECMs from protein behaviours that happen at the milliseconds *in vivo*. Establishing such predictive biophysical theory and spatiotemporal modelling system will allow computer simulations to replace some laboratory experiments to quantitatively test how new drugs and gene editing technologies targeting on proteins affect cell dynamics in the future.

### Spatiotemporal self-adaptive propulsion theory and multiscale modelling system

Branched actin network protrusion is an important way that drives cell migrations in ECMs. Through adding actin monomers to the barbed ends, polymerizing branched actin filaments grow and thus generate pushing force on the leading-edge membrane (5). Before being capped by capping proteins, the growing length of filaments with the polymerizing time *t* can be expressed as

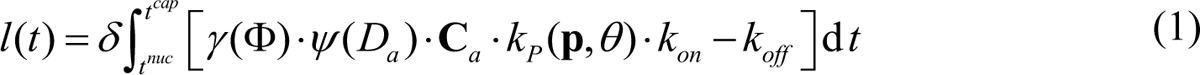

where *δ* is the radius of an actin monomer; *t^nuc^* and *t^cap^* are the nucleation time and the capping time, respectively; **C***_a_* is the local concentration of actin monomers in the cell; *γ* (Φ) is the consuming factor of actin monomers, introducing the relation that polymerizing rate is proportional to the ratio of the concentration of actin monomers to the density of polymerizing filaments _Φ_ (35); *ψ* is a scaled diffusion coefficient *D_a_* of actin monomers and is to introduce the effect of actin filament density on the actin diffusion flux towards the polymerizing barbed ends based on the Fick’s first law of diffusion; *k_P_* (**p**,*θ*) is an interacting force-induced probability density distribution in the rate of filament polymerization due to size-dependent insertion of monomers 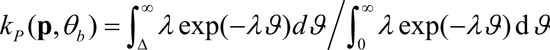(13, 36); **p** is the interaction force that the polymerizing actin filament acts on the leading-edge membrane; *θ_b_* is the angle of the filament barbed end relative to the normal vector of the local membrane; *ϑ* is a function of **p** and *θ_b_*, and Δ is the size of sufficient gap to permit intercalation of monomers(13, 36). *k_on_* and *k^off^* are the polymerization and depolymerization rate constants, respectively.

The leading-edge membrane under the polymerizing force of filaments is in a bending state. We simplify it into several continuous inclined planes based on the theory of differential geometry (Supplementary Fig. 2b). Though the mechanical interactions between all polymerizing actin filaments and the leading-edge membrane are in three-dimensional space, the interaction between a single polymerizing actin filament and the local membrane can be described in a two-dimensional deformation plane (Fig. 1b and Supplementary Fig. 2c). Then, based on the geometric nonlinear deformation theory of continuum mechanics, the spatial and temporal mechanical interactions between the growing (polymerizing) actin filament and the leading-edge membrane is derived (Fig. 1b, Supplementary Methods):

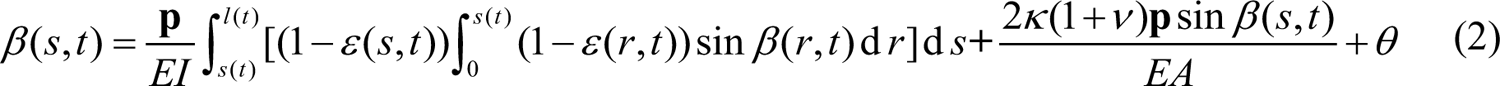

Where *β* (*s*, *t*) is the deformed angle along the actin filament due to the combined effects of bending, axial compression and transverse shear under the polymerizing growth; *E*, *ν*, *A*, *I*, *κ* are the Young’s modulus, Poisson’s ratio, cross-sectional area, the second moment of the cross-sectional area and shape factor of actin filaments, respectively. Using the deformation compatibility condition,

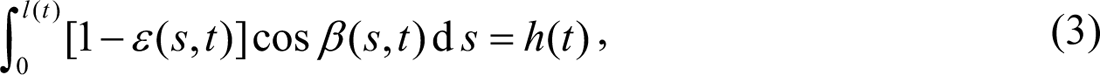

the nonlinear deformation function in Eq. (2) can be solved through iteration. *h* is the distance from the pointed end of the polymerizing actin filament to the local leading-edge membrane. Then, the interacting force **p**(*t*), the propulsive force **f***_p_*(*θ*,*t*) in cell migration direction, the total deformation energy *U* (*t*) and the mean bending curvature 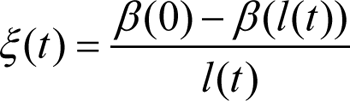 of the actin filament can all be solved (Supplementary Fig. 4a-f). The total deformation energy *U* (*t*) is expressed as

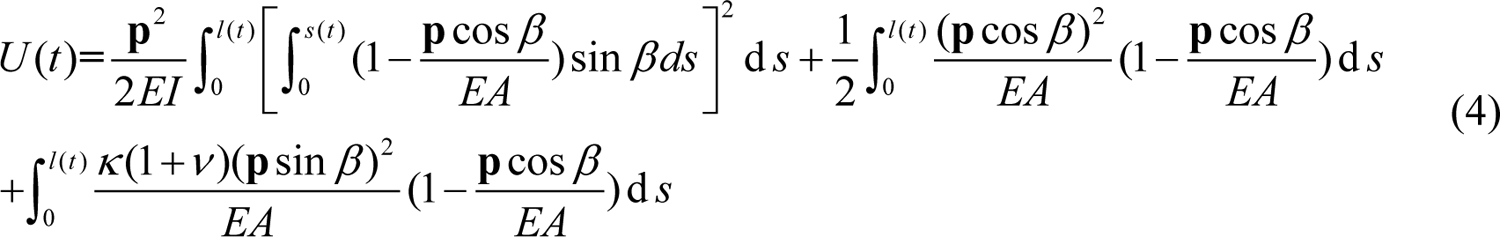

Our mechanical analysis also shows that convex side surface of the bending actin filament is stretched while the concave side surface is compressed (Fig. 1c). Their relative strain *ε_r_* = 2*r*_0_*ξ* where *r*_0_ is the radius of actin filament. Experiments(37) show that *in vitro* long actin filaments (∼10μm) exhibit large bending deformations under thermal fluctuations, and Arp2/3 complexes prefer to bind onto the convex sides of bending actin filaments. We fit the experimental data of relative branched density *P* with an inverted sigmoid function *P* = 2 (1+ *e*^0.73^*^ξ^*) (Fig. 1d) and analyse it. Strikingly, we have, for the first time, obtained the relative curvature-dependent Arp2/3-actin filament dissociation constant *K_d_*(**p**)*K_d_*^0^ = 0.5(1+ *e*^0.73^*^ξ^*) where *K_d_* (**p**) and *K_d_*^0^ are the dissociation constants in the bending and straight states, respectively (Fig. 1e and Supplementary information). The relative *K_d_* (**p**) *K_d_*^0^ shows that the affinity of Arp2/3 complex binding on the convex surface (negative curvature side) of actin filament is higher than the straight surface, which is higher than the concave surface (positive curvature side). This could be explained by the combination of our mechanical analysis and the recent cryo-electron structure of Arp2/3 complex-actin filament junction(38–40) (Fig. 1c and Supplementary Fig. 1). There are five actin subunits in mother actin filament that contact with Arp2/3 complex, with many of the contact surfaces in the grooves between these subunits (supplementary Fig. 1a)(38–40). ArpC1of Arp2/3 even has a protrusion helix that inserts into actin subdomains for binding (supplementary Fig. 1b)(38, 40). During the polymerization of actin filaments, they generate bending deformations under the constraints of cell membrane and ECM, leading to compressions on their concave sides (Fig. 1b and c). As a result, some of the binding surfaces in the grooves on the concave side is buried (Fig. 1c), inducing the decrease of the binding affinity of Arp2/3. On the contrary, the groove sites on the convex side of the bending actin filaments undergoes stretch (Fig. 1c), which facilitates Arp2/3 complex binding and improves Arp2/3 binding affinity. To better show this force-dependent binding affinity, we also calculate the relative dissociation constant *K*^1^_d_(**p**) / *K_d_*^2^(**p**) where *K_d_*^1^ (**p**) and *K_d_* ^2^ (**p**) are the dissociation constants on the convex and concave surfaces, respectively, demonstrating that the binding affinity on the convex surface is much higher than the concave surface (Fig. 1f). Our theoretical analysis has shown that although *in vivo* actin filaments are relatively short (∼250nm), their polymerization force still can induce them to generate significant backward bending deformations, which could be verified by the actin retrograde flow phenomenon (41) and the recent measurement that the bending curvature can reach up to 10 µm ^-1^ (42). This indicates that the force-dependent Arp2/3 complex-actin filament binding affinity occurs *in vivo*. To incorporate that Arp2/3 complex has a higher binding affinity with the convex surface of actin filaments than with straight surface, we introduce a bending curvature-dependent binding factor *d ^arp^* (*ξ* ^max^), which is defined as the space between two adjacent Arp2/3 complex branches along an actin filament, where *ξ* ^max^ is the biggest bending curvature in the deformation history of the actin filament (Eq. S14). The number of Arp2/3 complexes binding on the *i*th actin filament with a polymerization length *l_i_* (*t*) of can be determined as *n_i_^arp^* = *l*_i_ / *d^arp^* (*ξ* ^max^). The probabilities that an Arp2/3 complex binding on the convex and concave side of an actin filament are *P*(|*ξ*|) [*P*(|*ξ*|) + *P*(− |*ξ*|)] and *P*(− |*ξ*|) [*P*(|*ξ*|) + *P*(− |*ξ*|)], respectively. Thus, the total number of actin filaments *N*(*t*) pushing against the leading-edge membrane at time *t* is

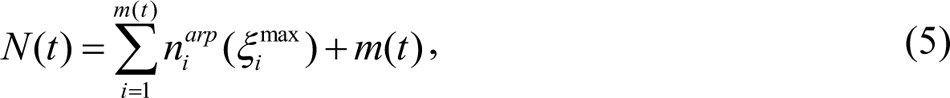

where *m*(*t*) is the number of mother filaments. During cell migrations, some filaments are linked to the leading-edge membrane through molecular linkers, such as Ezrin and N-WASPs(12, 43-45), and generate a resultant attachment force 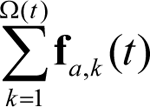 to pull back the membrane where Ω(*t*) and **f***_a_* are the total number of attaching molecular linkers and attaching force of each molecular linker, respectively. The total number of tethered filaments Ω(*t*) = *α N* (*t*), where the parameter *α* is the percentage of the total number of actin filaments contacting with the leading-edge membrane. In the static state, the leading edge of migrating cells follows force balance (Fig. 1g):

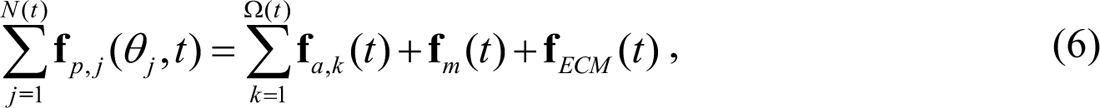

where **f***_m_* (*t*) and **f***_ECM_* (*t*) are a backward tension force from the leading-edge membrane and a resistance force from ECM, respectively. The total elastic deformation energy stored in all *N* (*t*) branched filaments pushing the leading-edge membrane is 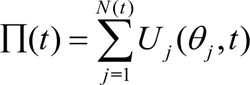. Experiments *j* **=**1 showed that the motion of branched actin filaments is saltatory with a step size Δ**S** of about 1-10nm with time (46, 47) due to the detachment of molecular linkers(13, 43, 48). Actually, in vivo, cell migration involves not only the detachments of molecular linkers from leading-edge membrane(43), but also the local ruptures of some nascent integrin adhesions (49) and extracellular crosslinking matrix networks owing to the propulsive force (21, 50). Since these processes are very complex and also involves different energy barriers, in order to capture the key characteristics of cell migration, we assume that when the resultant propulsive force 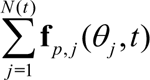 is larger than the maximum resultant stall force 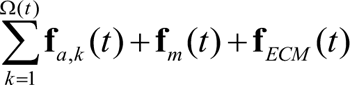, some *k* =1 molecular linkers will detach from the leading-edge membrane (43) and thus cell will migrate forward with a leaping step size Δ**S**. Then, the position of the leading edge of the cell at time *t* +1 can be expressed by

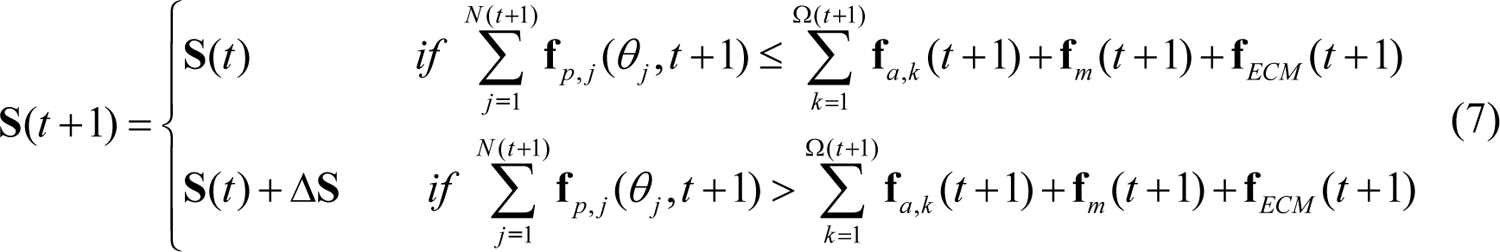

and the transient average migration velocity **V** is Δ**S** / Δ*t*. The dynamic leading-edge migration should meet the force balance defined by

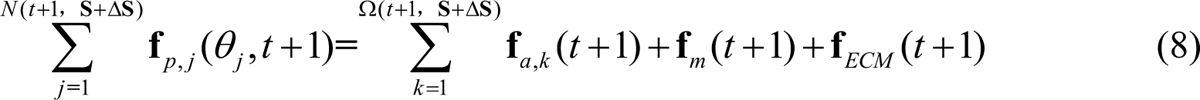

Because the present theory is derived from the spatiotemporal mechanical interactions between the polymerizing branched actin filaments and the leading-edge membrane constrained by extracellular resistance, and incorporates the force-dependent assembling behaviours of proteins, we name it spatiotemporal ‘resistance-adaptive propulsion’ (RAP) model.

To quantitatively shed light on how the assembling behaviours of multiple proteins impact on the dynamics of cell migration, we further develop a multiscale spatiotemporal modelling system by applying the RAP theory and integrating the stochastic behaviours of proteins (Supplementary Methods). This modelling framework systematically encompasses the *in vivo* actin monomer nucleation, actin filament polymerization, capping protein terminating filament polymerization, mechano-chemical nucleation of Arp2/3 complex, ATP binding, bent leading-edge membrane, detachments of molecular linkers, protein gradients caused by chemotactic cues, integrin-based adhesion and heterogeneous ECMs. Using this bottom-up approach, we can span multiple scales in both space and time to investigate cell migration behaviours in complex ECMs, and thus shed light on their dominating biophysical principles from the level of specific proteins.

### ECM resistance-triggered positive feedback adapts propulsive force

High extracellular resistance results in denser lamellipodial branched actin filaments during cell migrations(1, 5). To explore quantitatively whether our spatiotemporal RAP theory can reproduce this significant behaviour and reveal its underlying biophysical mechanism, we perform spatiotemporal simulations of cell migrations in both mechanically homogeneous extracellular microenvironment (HMECM) and HTECM (Fig. 2a). we observe that when the resistance **f***_ECM_* is 0.5 nN/µm (the normal range is 0.1–2.0 nN/μm of ECM(51)) in the HMECM condition, the density of polymerizing branched actin filaments Φ stably fluctuates in a very narrow range 230–270 /µm (Fig. 2a), agreeing well with the experimental data 150–350 /μm(51). However, in the HTECM, when the extracellular resistance **f***_ECM_* increases from the low resistance (LR) 0.5 nN/µm to a higher resistance (HR) 1.0 nN/µm, the branched actin density Φ increases by ∼35% (320–350 /µm, Fig. 2a). After **f***_ECM_* returning to the low resistance (RLR) 0.5 nN/µm, the density also decreases to its previous level (Fig. 2a). The density of Arp2/3 complex assembling actin cytoskeleton cofluctuates with the resistance (Fig. 2b). Strikingly, our model well predicts the experimentally measured correlations between the extracellular resistance and the lamellipodial branched actin filament density during cell migration in refs.(1, 5, 11) (Fig. 2a). The architecture of the lamellipodial branched actin network generated in our spatiotemporal simulations is also consistent with the experimental measurements (Fig. 2c) (52). In addition, we show that when cells encounter higher resistances, the consumption rate of ATP (Fig. 2d) increases to fuel cell migrations. This prediction is also validated by the experimental data (Fig. 2d) that mitochondria and ATP levels are higher at the invasive cell leading edge in a stiffer ECM confinement(11) and that cancer cells overproduce ATP to boost their lamellipodia formations and invasions(53).

**Fig. 2.**
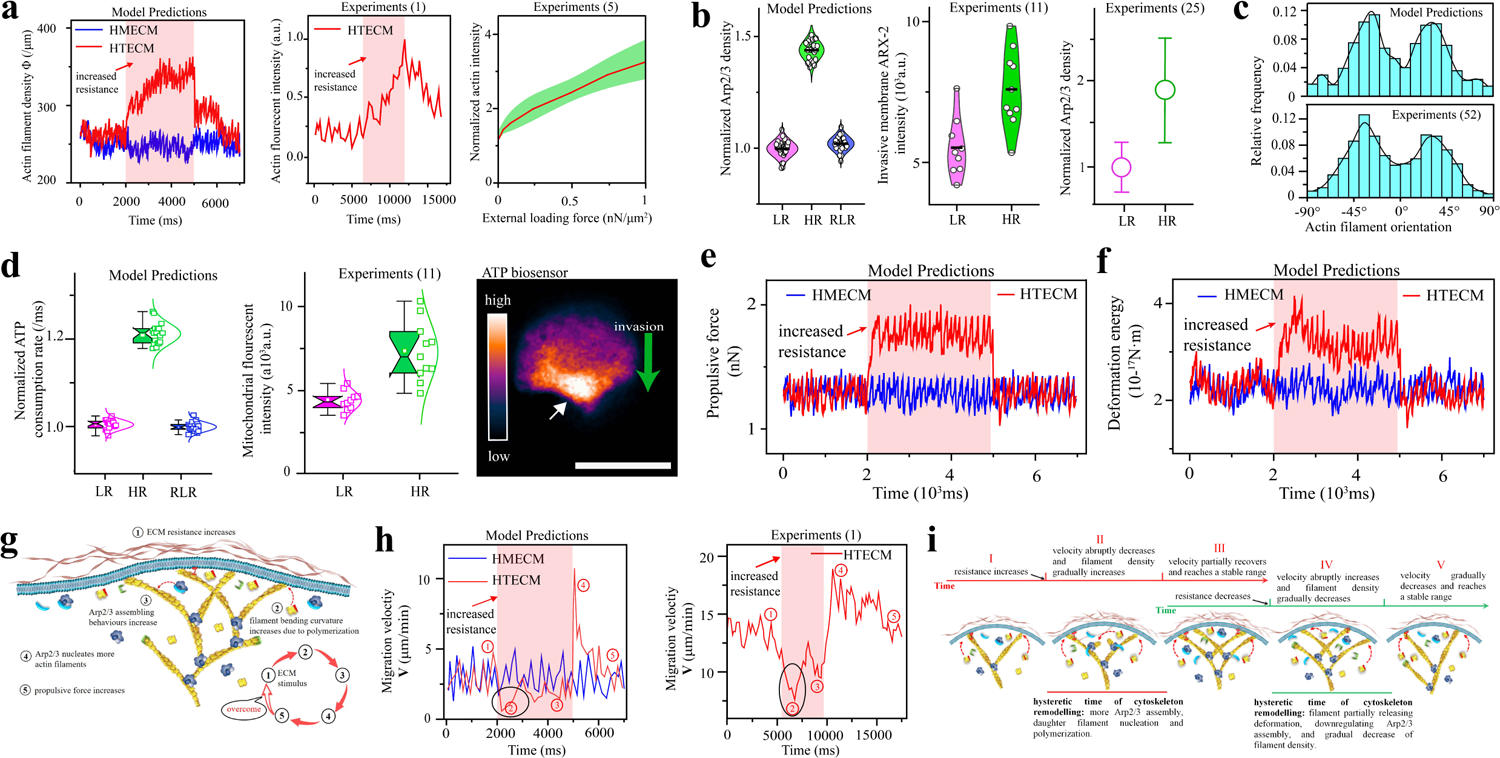
Spatiotemporal simulations from protein behaviours accurately predict self-adaptive cell migrations in complex ECMs and reveal their underlying biophysical mechanisms. (**a**) Resistance force-induced branched actin network density adaption. Model predictions: temporal fluctuations of polymerizing actin cytoskeletal density at the leading edge in HMECM (blue line) and HTECM (red line). The extracellular resistance f*_ECM_* of the HMECM is 0.5 nN/μm. For the HTECM condition (red line), the ^f^*ECM* is 1.0nN/μm in the time frame 2000–5000 ms (shadow region), and is 0.5 nN/μm in the time frames of 0–2000 ms and 5000–7000 ms. The induced actin density along the leading edge in the two conditions is 200–350 /μm, agreeing well with the experimental data 150–350 /μm(51). The first experimental data is the temporal fluctuations of actin reporter lifeact: GFP intensity at the leading edge of migrating fish keratocytes in HTECMs from experiments in ref. (1). The second experimental data is the fluctuations of polymerizing branched actin density in responding to varying external loads from experiments in ref.(5). Actin density is normalized by that of the unloaded condition(5). Green region is the standard deviation. (**b**) Resistance force-induced Arp2/3 complex density adaption. Model predictions: temporal fluctuations of the leading-edge Arp2/3 density in responding to HTECMs (n=30 each group). The Arp2/3 density is normalized by that of the homogeneous ECM condition. The first experimental data: adaptive fluorescent intensity of ARX-2::GFP for Arp2/3 complex at the leading edge when cells invade ECMs with low resistance and high resistance (n=10 each group) from experiments in ref. (11). The second experimental data: Arp2/3 complex density under increasing resistance (n=15 each group) from experiments in ref. (25). The HR condition is normalized by the LR condition. (**c**) Model predictions and published experimental measurements(52) of the orientation frequency of lamellipodial actin filaments relative to the migration direction, which is defined as 0 ° direction. (**d**) Adaptive ATP consumption rate for fuelling cells migrating through HTECMs. Model predictions: the ATP consumption rate is the average number of ATP used for assembling actin cytoskeleton per micro-second, and is normalized by that of the HMECM condition (n=20 each group). Experiments: adaptive fluorescent intensity of mitochondria and ATP at the leading edge when cells invade ECMs with low resistance and high resistance (n=10 each group) from ref. (11). Scale bar is 5 μm. Temporal fluctuations of propulsive force (**e**) and deformation energy (**f**) generated by the leading-edge actin cytoskeleton in HMECM and HTECM. (**g**) Molecular protein-level biophysical mechanism of the positive feedback that migrating cells mechano-sense the variation of ECM and then accurately adapt their propulsive forces to overcome it. (**h**) Velocity adaptation. Model predictions: temporal fluctuations of leading-edge migration velocities in HMECM and HTECM. Experiments: temporal fluctuations of leading-edge migration velocity in responding to HTECM from the experiments in ref.(1). (**i**) Molecular protein-level biophysical mechanism of leading-edge migration velocity depending on extracellular resistance history is the temporal hysteresis of the leading-edge actin cytoskeleton remodelling and adaptation. The red and green time lines correspond to the velocity adaptations caused by the increased and decreased ECM resistances, respectively. Velocity adaptation stages Ⅰ–Ⅴ in (**i**) correspond to the stages Ⅰ–Ⅴ in (**h**), respectively.

Next, we analyse the spatiotemporal propulsive force (Fig. 2e) and the elastic deformation energy _Π_ stored in the branched filaments (Fig. 2f). It is found that both synchronously fluctuate with the density of branched actin filaments. We further quantitatively identify that the adaptation of filament density is to meet the propulsive force and energy demands for overcoming the varying extracellular resistance. To shed light on the more fundamental cross-scale biophysical mechanism of such significant adaptive behaviours in response to ECMs, we then examine the assembly of protein molecules that happens at milliseconds, finding an ECM resistance-triggered positive feedback (Fig. 2g). When the resistance **f***_ECM_* increases, polymerizing (growing) actin filaments under the ECM confinement will automatically have larger nonlinear bending deformations (Supplementary Movie), which increase the probability that Arp2/3 complex will bind and nucleate daughter actin filaments, and vice versa. Through this mechanism, cells can adapt the density of branched actin network and thus the propulsive force and energy for migrations. More importantly, through the extent of bending deformations, cells can sense the ECM resistance, so a larger resistance induces a larger bending deformation. Thus, from the protein level, we reveal that migrating cells can sensitively sense the immediate variations of ECM resistance through the polymerizing growth of actin filaments, and then accordingly make accurate adaptive responses in filament density, propulsive force and energy through the mechano-triggered Arp2/3 complex-actin filament assembling behaviours (Fig. 2g). This mechanism also endows migrating cells with an optimization ability to accurately employ their intracellular proteins and ATP (Fig. 2a, b and d) according to their demands in complex ECMs.

### Leading-edge velocity depends on the temporal hysteresis of filament density adaptation to varying ECM

We next explore how the leading-edge migration velocity responds to the varying stiffness of the ECMs and how the underlying biophysical mechanisms operate at the level of protein molecules. The spatiotemporal simulations in HTECMs show that when migrating cell encounters an increased extracellular resistance **f***_ECM_*, its leading-edge migrating velocity **V** suddenly decreases from 3.3 µm/min to 0.7 µm/min (from stage Ⅰ to Ⅱ in the black ellipse in Fig. 2h), which is lower than the velocity 1.7–5.2 µm/min in the HMECMs with **f***_ECM_* = 0.5 nN/µm. However, with the continuing polymerization of branched filaments, the decreased velocity partially recovers. This is owing to the gradual increase in the filament density based on the ECM resistance-triggered positive feedback. Afterwards, we reduce the extracellular resistance from 1 nN/µm to its previous value 0.5 nN/µm for the HTECMs. Strikingly, the leading-edge velocity abruptly increases to a very high value 10.7 µm/min, and then gradually decreases to the previous range 1.7-5.2 µm/min (Fig. 2h). The spatiotemporal predictions from stage Ⅰ to Ⅴ are validated by the experimental data (Fig. 2h) (1, 26). To gain insight into the protein behaviours, we check the evolution of the spatiotemporal remodelling of the growing lamellipodial actin network. Previous studies(26) described this phenomenon as velocity dependence on loading history. Here, we quantitatively show that the nature of the leading-edge velocity variations in the HTEMCs stems from that the adaptation of branched actin filament density is always temporally hysteretic to its triggering reason, i.e., varying ECM resistance (Fig. 2i). This is because the generation of daughter filaments and their growths to the leading-edge membrane always cost some time due to actin monomer nucleation, filament polymerization, Arp2/3 activation and assembly (Fig. 2i). Although the extracellular resistance **f***_ECM_* has increased or decreased, the density, propulsive force and deformation energy of polymerizing actin filaments keep unchanged in this process. Thus, the velocity suddenly decreases because of incapable of overcoming the increased resistance, and increases because of easily overcoming the decreased resistance to release the excess deformation energy, respectively (Fig. 2e and f). This also indicates that, in HTECMs, there exists no one-to-one correspondence between **V-f***_ECM_* that can describe the leading-edge migrating behaviours. However, in HMECMs, the one-to-one **V-f***_ECM_* relationship exists.

### High ECM resistance-triggered negative feedback adapts cell morphology for pathfinding

The above study focuses on the low ranges of ECM resistance. However, the leading edge usually faces some local dense collagen regions with high resistances in ECM. Thus, we here investigate migratory pathfinding in this kind of complex ECMs and shed light on its underlying mechanistic operating basis at the protein level. We design an ECM as demonstrated by Fig. 3a. The ECM is divided into two stages. In the first stage (from 0 to 110 nm), it is mechanically homogeneous and has a resistance **f***_ECM_* of 0.5 nN/µm. However, in the second stage (from 110 to 300 nm), its resistance becomes 1.0 nN/µm, and there are two very dense collagen regions with a very high resistance **f***_ECM_^right^* = **f***_ECM_^left^* = 5 nN/µm.

**Fig. 3.**
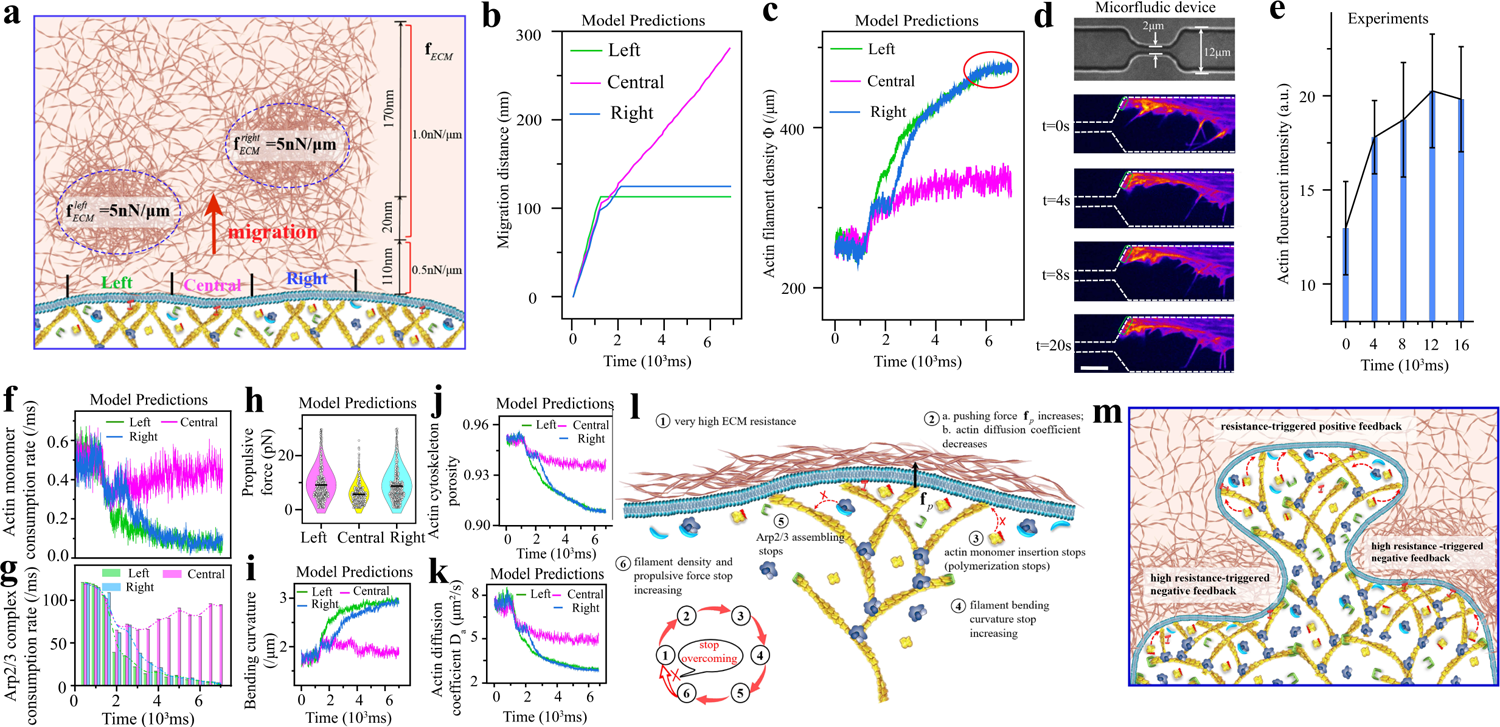
High extracellular resistance triggers negative feedback to adapt cell morphology to circumnavigate obstacles for compromisingly pathfinding. (**a**) The designed HTECM in simulations. It is divided into two stages, where extracellular resistances in the first (in the range of 0–110 nm in front of the migrating cell leading edge) and second stages (in the range of 110–300 nm in front of the migrating cell leading edge) are 0.5 nN/µm and 1.0 nN/µm, respectively. However, the second stage has two dense ECM regions with a very high resistance f*^left^* = f*^right^* = 5 nN/µm. The red arrow denotes cell migration direction. Spatiotemporal migration distances (**b**) and leading-edge actin filament densities (**c**) at the left, central and right parts of the leading edge. (**d** and **e**) Microfluidic device and temporal fluctuation of actin reporter GFP-lifeact: the actin intensity as a function of the time at the leading edge when migrating cells encounter an obstacle (n=10 each group). Scale bar in (**d**) is 5 µm. Temporal actin monomer (**f**) and Arp2/3 complex (**g**) consumption rates at the left, central and right local leading edges. Actin monomer consumption rate is the ratio of the number of actin monomers adding to the barbed ends of actin filaments to the number of uncapped actin filaments per micro-second. Arp2/3 complex consumption rate is the number of Arp2/3 complex assembling for the actin cytoskeleton in a time span of 500 ms. (**h**) Propulsive forces produced by each polymerizing actin filaments at the left, central and right local leading edges. Temporal fluctuations of the average bending curvature of polymerizing actin filaments(**i**), the leading-edge actin cytoskeleton porosity (**j**) and the actin diffusion coefficient (**k**). (**l**) Biophysical mechanism of the high ECM resistance-triggered negative adaptation feedback. (**m**) Demonstration of simulation results that the leading edge circumnavigates high ECM resistance regions based on the negative feedback and opens a channel in the weak region based on the positive feedback. The positive and negative feedbacks work cooperatively to adapt cell morphology and drive cell migration.

Spatiotemporal simulations show that, in the first homogeneous mechanical environment, the left, central and right parts of the leading edge migrate forward synchronously with similar velocities (Fig. 3b). The density of branched actin filaments is approximately homogeneous (Fig. 3c). However, when the cell encounters the two dense collagen ECM regions, it stops moving forward from the left and right sides, and turns to squeeze out from the central region where the resistance is weak (Fig. 3b). Unexpectedly, even though the leading edges on the left and right sides have not overcome their local resistances, the increase of the density of actin filaments stops (red ellipse area in Fig. 3c). Interestingly, this indicates that when the ECM resistance is very high, the mechanism of resistance-triggered positive feedback no longer work. We perform experiments to validate this spatiotemporal model prediction. We record Human retinal pigment epithelial-1 (RPE1) cells expressing green fluorescent protein linked to a small peptide (Lifeact-GFP) with affinity for actin microfilaments. We image the dynamic cell migration processes in the microchannels with constrictions (Fig. 3d), and track the temporal actin intensity when the cell encounters the constriction regions. The experimental results of the temporal variations of leading-edge actin intensity are strongly consistent with our modelling predictions (Fig. 3e). We then check how our spatiotemporal simulations operate at the protein level (Fig. 3f and g). Migrating cells apply the resistance-triggered positive feedback to adapt filament density and propulsive force to try to overcome the obstacles until high forces between the leading-edge membrane and actin filaments (Fig. 3h) compromise the intercalations of actin monomers with the barbed ends (Fig. 3f), which stops the bending deformation of filaments (Fig. 3i) and the assembly of Arp2/3 complex (Fig. 3g). These results of our simulation are strongly validated by *in vitro* experiments (25). We also find that the porosity of the leading-edge actin network is reduced due to the increased density of actin filaments (Fig. 3j). This lowers the diffusion of actin monomers to the free barbed ends, and reduces the polymerization rate at the leading edge (Fig. 3k).

Unexpectedly, we find that an ECM resistance-triggered negative adaptation feedback (Fig. 3l) coexists with the resistance-triggered positive adaptation feedback (Fig. 2g) in migrating cells. While the positive feedback adapts cell propulsive force to overcome ECMs, the negative feedback can adapt cell morphology by stopping actin polymerization to circumnavigate the high resistance regions in ECMs. This behaviour can avoid unnecessary consumptions of intracellular proteins and ATP resources (Fig. 3f and g), and thus improve cell migration efficiency. The synergy of the two opposite feedbacks allow the leading edge to discriminate whether the local ECM confinement is weak to overcome or strong enough to require circumnavigation with a formidable efficiency. It endows cells with both powerful and flexible migration capabilities to widely adapt morphologies to the complex ECMs (Fig. 3m). This may explain why cancer metastasis is extremely hard to prevent once they acquire invasive ability. In addition, our results reveal that the initiation of cellular morphology adaptations, a prominent characteristic of invasive cancer cells, derives from the leading edge sensing strong barriers and escaping from them.

### Directional cell migration is steered by a balanced and competing relation

Cells typically follow the gradients of chemotactic cues to migrate(6). The nucleus of migrating cells acts as a mechanical gauge to make a temporary adaptations while choosing a passable path (6, 54, 55). Given the global migration direction predefined by chemotactic cues, leading edge is much more important in persistently directing cell migration towards the final destination. Our simulations demonstrate that different local stiffness of ECM results in different local densities of branched filaments along the leading edge. Here, we investigate another challenging problem of how the complex interplays between the multiple simultaneous factors of extracellular resistance, density heterogeneity of branched actin filaments, and the external diffusible chemotactic stimuli synergistically steer directional cell migration in HTECM. We introduce a gradient diffusion of a localised chemotactic cue sensed by transmembrane receptors (Fig. 4a), rendering a gradient distribution of intracellular actin monomers. We design and simulate three cases A–C (Fig. 4b), in which local leading edges at the positions of **x**_1_ and **x**_2_ simultaneously drive cell migrations. Based on the experimental observations (6), it is reasonable to hypothesize that the fastest migration direction of local leading edges is the main migration direction of a cell. Our simulations in case A show that the migration distance of the leading edge at the position **x**_2_ are larger than those at the position **x**_1_ (Fig. 4c), concluding that when the extracellular mechanical microenvironments are homogeneous, cells more actively migrate toward the site where the local concentration of actin monomers is higher. Since homogeneous ECM induces homogeneous density of branch filaments in the whole leading edge (Fig. 4d), a higher local concentration of actin monomers means that sufficient actin monomers can be supplied to the polymerizing barbed ends of the local branched actin filaments. The chemotactic cue determines the local protein concentration to steer directional cell migration. Then, we test case B, in which the left and right sides are designed to have the same local concentrations of actin monomers due to homogeneous distribution of the chemotactic cue, but different extracellular resistances. Cells migrate toward the low resistance **x**_1_ side (Fig. 4e), because higher resistance on the right-side results in denser branched actin filaments there (Fig. 4f), which deplete action monomers locally and thus slows polymerization rate at barbed ends. This case highlights that extracellular resistance plays a determining role in directing cell migration. However, the results of case C (Supplementary Fig. 5), in contrary to case B, shows that cells migrate toward the high resistance **x**_2_ side (Fig. 4g). Even though the denser polymerizing filaments consume more action monomers on the **x**_2_ side (Fig. 4h), the strong chemotactic cue can sustain a high local concentration of actin monomers, which enable branched actin filaments to polymerize at a higher rate than that on the **x**_1_ side. Although here we take actin monomers as an example, local variations in the concentrations of other intracellular proteins, such as Arp2/3 complex, WASPs, FMNL, aprin and profilin, have the same effect on cell migration by enhancing or inhibiting nucleation, branch formation and filament polymerization.

**Fig. 4.**
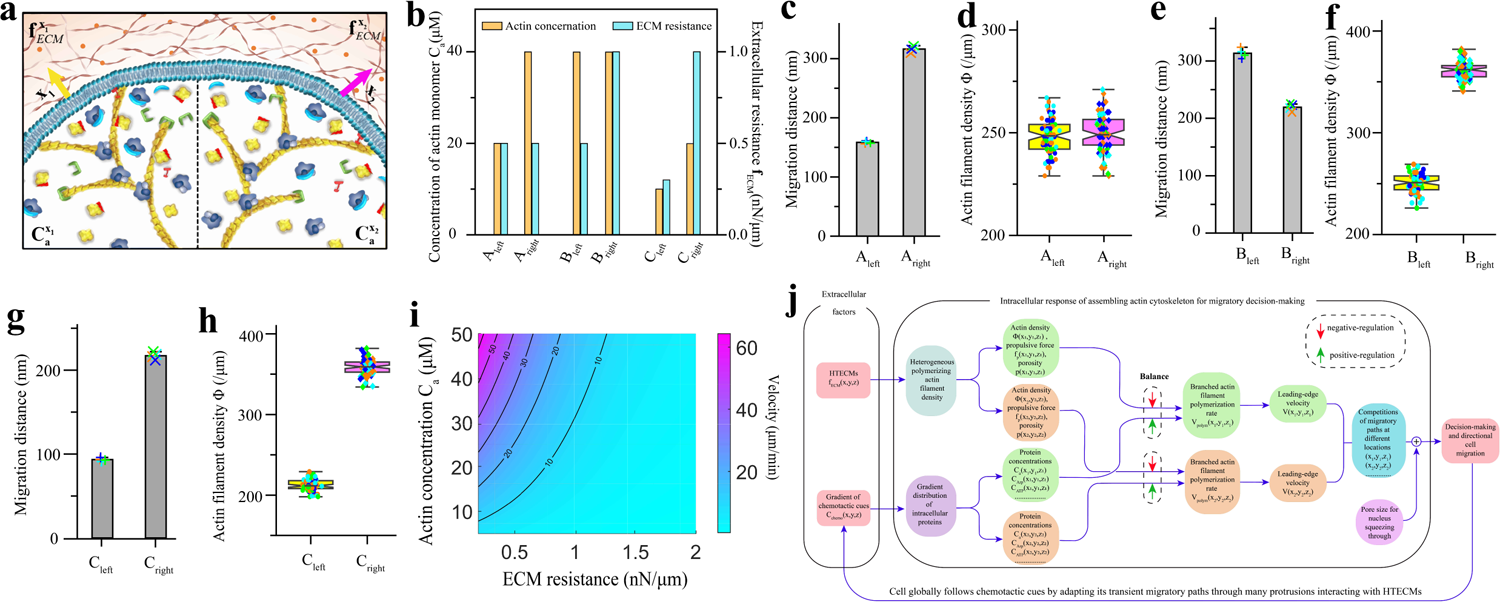
Directional cell migration is steered by a balanced relation between intracellular proteins, HTECMs and chemotactic cues. (**a**) Demonstration of initial simulation conditions that local leading edges drive cell migration at the two locations of **x**1 and **x**2. Yellow and violet arrows denote the migration directions at **x**1 and **x**2, respectively. The dots represent chemoattractant of actin. (**b**) Local extracellular resistance of ECMs and local concentration of actin monomers caused by a gradient chemoattractant at the positions of **x**1 and **x**2 in cases A-C. Migration distances (**c**, **e** and **g**) and leading-edge actin cytoskeleton densities (**d**, **f** and **h**) in cases A-C (n=4). (**i**) Leading-edge migration velocity for varying ECM resistance and actin concentration. (**j**) Biophysical mechanism of directional cell migration is that it is steered by a balanced relation between intracellular proteins for assembling leading-edge actin cytoskeleton, HTECMs and chemotactic cues.

From the distinct results of cases A–C, we find that directional cell migration is not solely steered by either the gradients of intracellular proteins induced by chemotactic cues or the local stiffness of ECMs. It is a balanced competing consequence between the two (Fig. 4i and j). While strong chemotactic cues improve the local nucleation, branch formation and polymerization rate of actin filaments, stiffer local ECMs result in denser local branched actin filaments, which in turn reduce these rates and slow down cell migration. In addition, through the collaboration of the resistance-triggered positive feedback mechanism and the high resistance-triggered negative feedback mechanism, here our results further indicate that as long as the chemotactic cues are strong enough in sustaining the high gradients of proteins for keeping a high branching and polymerizing rates, the leading edge will drive cells to migrate globally and persistently toward the prescribed final destination by overcoming weak confinements and circumnavigating strong barriers encountered.

## Discussions

Although cell migrations have been studied many decades, how cells mechano-sense and make self-adaptive responses to complex ECMs at the protein level still remains elusive. In this study, by analysing the spatiotemporal nonlinear deformation of polymerizing actin filaments and the polymerization force-regulated Arp2/3 complex-actin filament binding interactions, we derive a spatiotemporal ‘resistance-adaptive propulsion’ (RAP) theory for cell migrations. Then, with the RAP theory, we develop a spatiotemporal multiscale modelling system, which can not only simulate dynamic cell migrations in ECMs, but also shed light on the assembling behaviours of single proteins. Our simulations predict many important spatial and temporal adaptive cell migration behaviours observed experimentally and clinically (Table 1), and reveal their underlying operating mechanisms emerged from proteins behaviours.

**Table 1.**
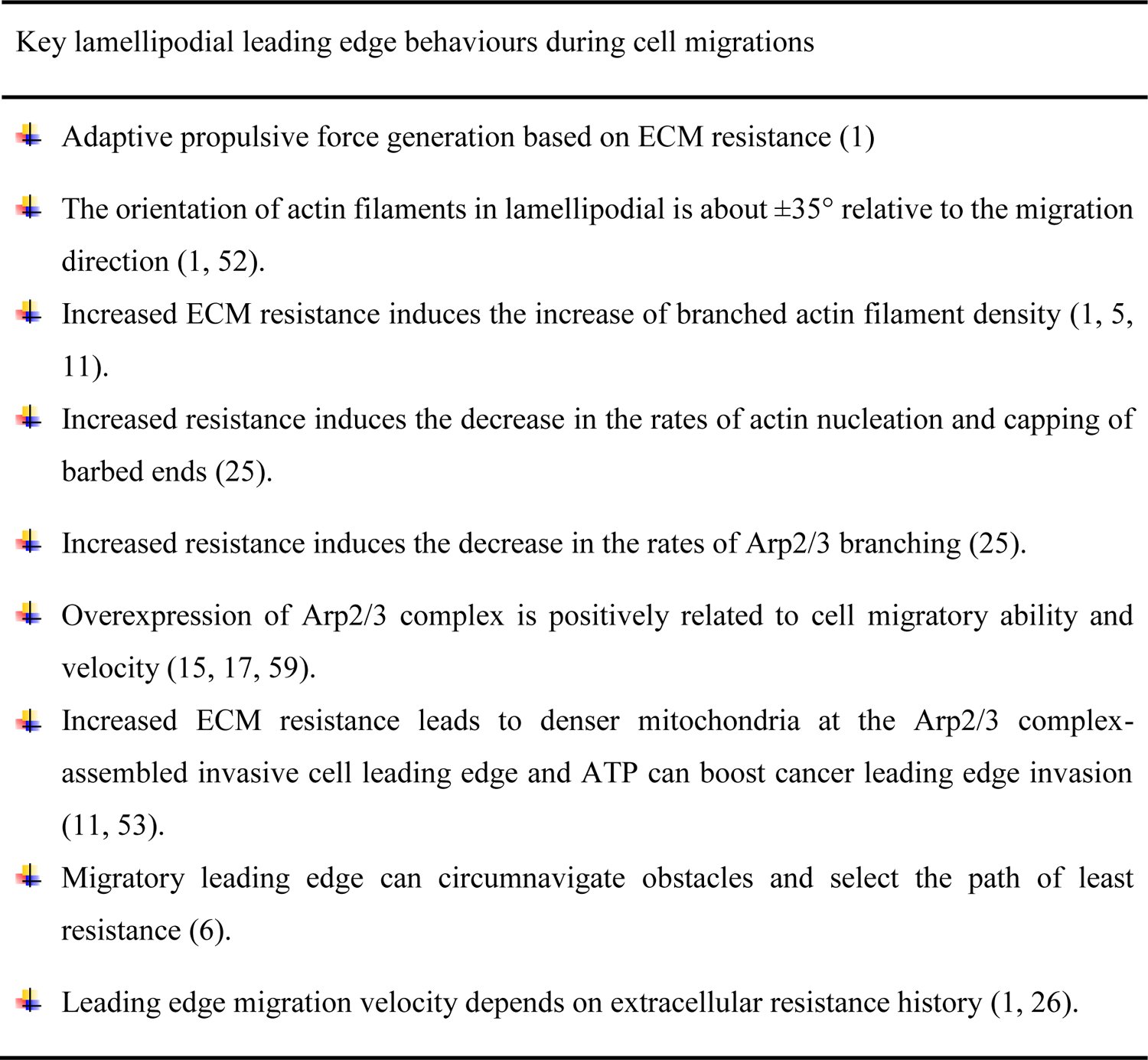
RAP model systematically predicts all the key adaptive migratory behaviours discovered in experiments.

We find that it is the polymerization force-regulated actin filament-Arp2/3 complex binding interaction that dominates self-adaptive cell migrations in complex ECMs, through the synergistic effects of positive and negative feedbacks. The polymerization-induced bending state of actin filaments serves as the mechano-sensor of cells. Our finding is highly consistent with two recent studies that show bending force (42) and polymerization force (56) evoke nucleotide-state conformational changes of actin filaments and thus affect the binding behaviours of actin-binding proteins. In fact, mechanical force-induced conformational change of proteins is a key regulator of protein-protein interactions (57), thereby regulating physiological and pathophysiological cellular behaviours.

Previous studies indicated that increased extracellular resistance induced denser branched actin filaments in the migration leading edge(1, 5, 25). Here, we identify that the enhanced leading-edge actin filament density is to improve the propulsive force and energy to overcome the resistance. Our results physically interpret why Arp2/3 complex overexpression is tightly associated with cancer cell invasions(15–19), poor patient survival in cancers(58) and high migratory force(59). We indicate that Arp2/3 is a key target protein for developing anti-cancer drugs, especially for patients in the advanced stages of cancer. It should be mentioned that the current simulations are for the conditions that integrin-based adhesions are sufficient for fixing the branched actin network. In the event that the levels of integrin or vinculin are not abundant, it is conceivable that high ECM resistance could trigger a rapid actin retrograde flow (41, 60) based on the positive feedback, resulting in an increased backward movement of the branched actin network and subsequently reducing the velocity of cell protrusion. This represents an additional type of negative feedback mechanism that can arise from excessively high ECM resistance.

It should be emphasized that polymerization of Arp2/3 complex-branched actin filament network is the most important way for cells to generate propulsive forces to interact with their surrounding microenvironments and perform their functions. It participates in phagocytosis of immune responses(61), endocytosis(62), dendritic spine formation of cortical neurons(63), T-cell and cancer cell interactions(64), and promoting of DNA fork repair (65) and chromatin organization(66, 67). All these processes require cells to mechano-sense the surrounding microenvironments and then make adaptive force generation responses. Thus, besides cell migrations, our cross-scale findings and the spatiotemporal multiscale modelling system can also be applied to investigate these dynamic physiological cell activities.

## Materials and Methods

### Cell culture

Human telomerase-immortalized, retinal-pigmented epithelial cells expressing GFP-LifeAct (CellLight Actin-GFP, BacMam 2.0 from Thermo Fischer Scientific) were grown in a humidified incubator at 37°C an d 5% CO_2_ in DMEM/F12 supplemented with 10% fetal bovine serum and 1% penicillin-streptomycin. All cell culture products were purchased from GIBCO/Life Technologies. Cell lines were regularly checked for mycoplasma contamination (MycoAlert, Lonza).

### Cell fixation and immunostaining

Cells were pre-permeabilized in 0.5% Triton X-100 in cytoskeleton buffer for 15 s for tubulin and then fixed in 0.5% glutaraldehyde (no.00216-30; Polysciences) in cytoskeleton buffer with 0.5% Triton X-100 and 10% sucrose for 15 min at room temperature. Cells were then washed three times with PBS-tween 0.1% and incubated in a quenching agent of 1mg ml-1 sodium borohydride for 10 min at room temperature. After fixation, the cells were washed with PBS-Tween 0.1% and then blocked with 3% bovine serum albumin (BSA) overnight. The cells were incubated with appropriate dilutions of primary antibodies in PBS containing 3% BSA and 0.1% Tween overnight at 4 °C in a humid chamber. After washing three times with PBS-tween 0.1%, the coverslips were then incubated with appropriate dilutions of secondary antibodies diluted in PBS containing 3% BSA and 0.1% for 1 h at room temperature in a humid chamber. After washing three times with PBS-Tween 0.1%, coverslips were then mounted onto slides using Prolong Gold antifade reagent (no. P36935; Invitrogen). Here, we used rat monoclonal antibodies against α-tubulin (no. ab6160, Abcam) and Alexa Fluor 555 goat anti-rat (1:500, no.A21434; Invitrogen) as secondary antibody. Antibodies were diluted as followed for immunofluorescence: α-tubulin (primary: 1:500) and secondary (1:1000). For actin immunofluorescence, Phalloidin-Atto 488 (49409, Sigma-Aldrich) was used to and diluted as 1:1000 to stabilize actin filaments in cells.

### Cell migration under constriction

Micro-channels were prepared as previously described (doi:10.1007/978-1-61779-207-6_28). Briefly, polydimethylsiloxane (PDMS, 10/1 w/w PDMS A / crosslinker B) (GE Silicones) was used to prepare 12 μm wide and 5μm high micro-channels with a constriction of 2 μm. For confined migration in Fig. 3d, coverslip and micro-channels were treated by plasma for 1mins and then were stuck together at 70 °C for 10mins. Before cell seeding, the microchannels were treated with fibronectin at 10 μg ml-1 for 30mins and incubate with culture medium for 3 hrs at room temperature. GFP-LifeAct RPE1 cells were then seeded in the microchannels with a concentration at 2×105 cells ml-1. Imaging was performed after the overnight incubation of the microchannels.

### Imaging

Images of the immunostainings were acquired on a Zeiss LSM900 confocal microscopes (Axio Observer) using a 63x magnification objective (Plan-Apochromat 63X/1.4 oil). Image acquisition for time-lapse of GFP life-act RPE1 cells in constrictions was performed on a confocal spinning-disc system (EclipseTi-E Nikon inverted microscope equipped with a CSUX1-A1 Yokogawa confocal head, an Evolve EMCCD camera from Roper Scientific, Princeton Instruments) through an 63x magnification objective (Nikon CFI Plan Fluor 60X/0.7 oil) objective every 4 s during 2 hrs for each time-lapse. The set-up was equipped with a live cell chamber, and the temperature was constantly kept at 37 °C. Labelled -actin was excited with a 491 nm laser line, and emission was observed with a standard GFP filter. The microscope was monitored with MetaMorph software (Universal Imaging).

### Spatiotemporal theoretical model and dynamic multiscale modelling system

The derivation of the spatiotemporal biophysical theory and the developing process of the spatiotemporal multiscale dynamic modelling system are provided in the Supplementary Text.

## Data availability

The data that support the findings of this study are available from the corresponding author on reasonable request.

## Author contributions

X.D.C., X.Q.F., H.X.Z. and M.G. designed the research. X.D.C., H.X.Z. and X.Q.F. developed the theory and the spatiotemporal simulation framework. X.D.C performed the simulations. X.D.C., B.W.X. and Y.H.M analysed the data. X.D.C. and Y.H.L. did the experiments and analysed experimental data. X.D.C., X.Q.F., H.X.Z., Y.H.L. and M.G. wrote the manuscript.

## Acknowledgements

We gratefully thank Professor Thomas D. Pollard for his invaluable insights and great help for manuscript preparation, and Professor Laurent Blanchoin for help with the experiments. We also thank Professors Xin Liang, Congying Wu, Bo Li and Yue Shao for valuable discussions. X.D.C., X.Q.F, and B.W.X. acknowledge the support from the National Natural Science Foundation of China (Grant no. 11921002 and 12032014)

## Code availability

All computer codes are available from the corresponding authors on reasonable request.

---

**Supplementary Fig. 1.**
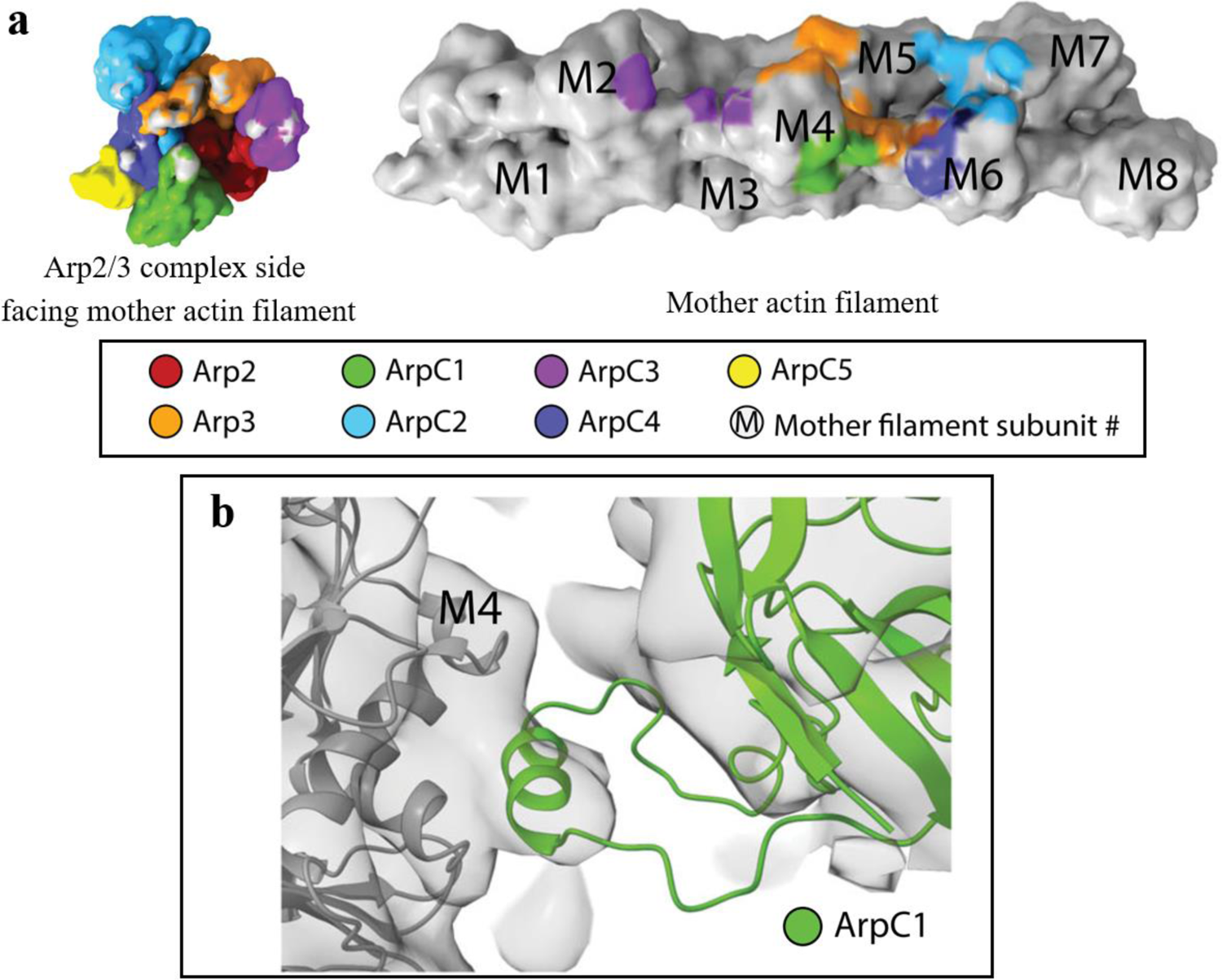
Actin-Arp2/3 complex binding interaction surfaces within the branch junction (38). (**a**) Binding interaction surface between mother actin filament and Arp2/3 complex. Most of the binding surface locates in the groove between actin subunits of the mother actin filaments. (**b**) The protrusion helix of ArpC1 of Arp2/3 complex insert into the mother actin filament subunit M4.

**Supplementary Fig. 2.**
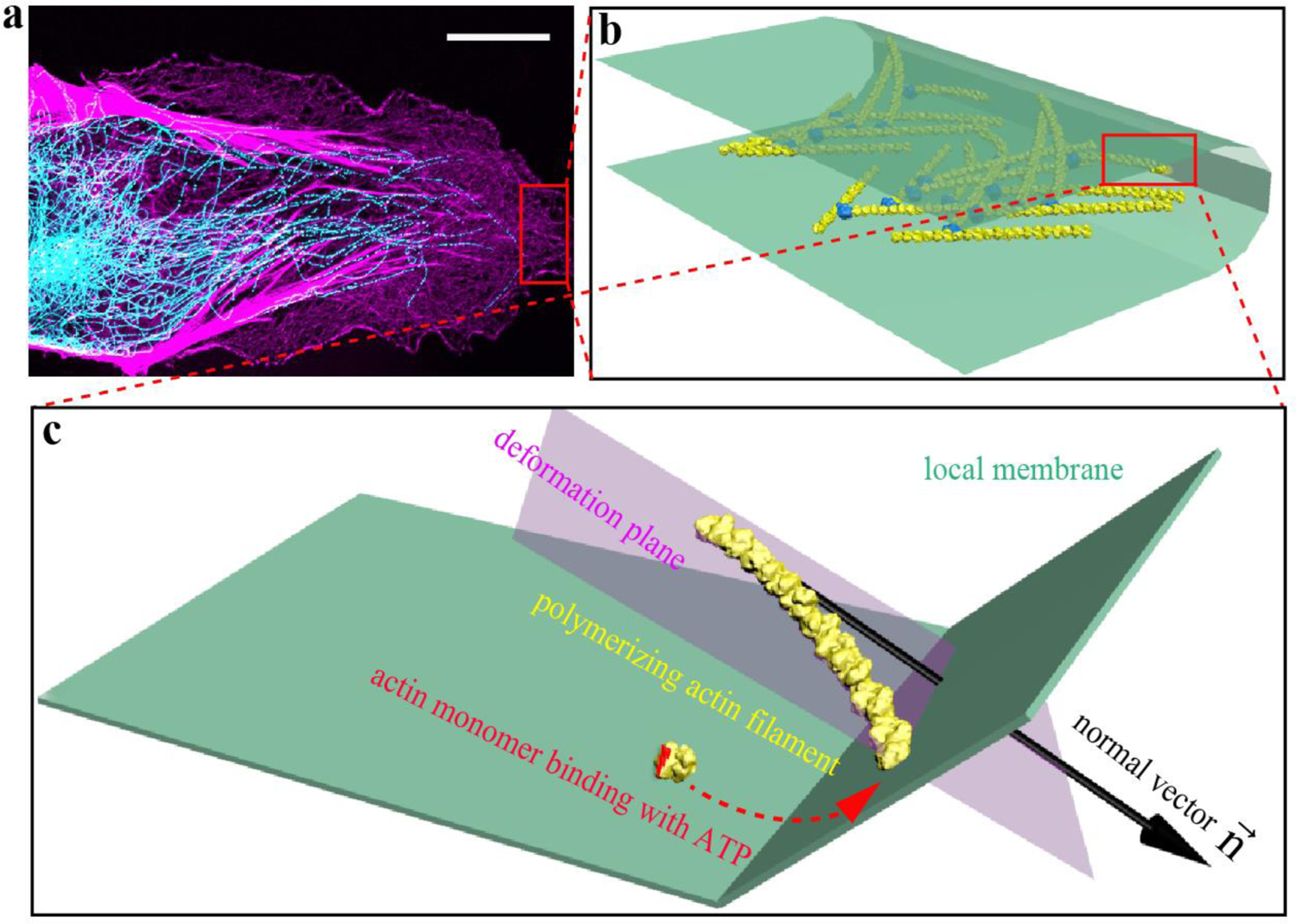
Polymerizing branched actin network drives cell migration. (**a**) Confocal image of a labelled migrating cell (actin: violet color; microtubule: cyan color); scale bar: 10 µm. (**b**) Polymerizing branched actin filaments under the leading-edge membrane. (**c**) The interaction between a single polymerizing branched actin filament and the local leading-edge membrane confined by ECM is a two-dimensional behaviour.

**Supplementary Fig. 3.**
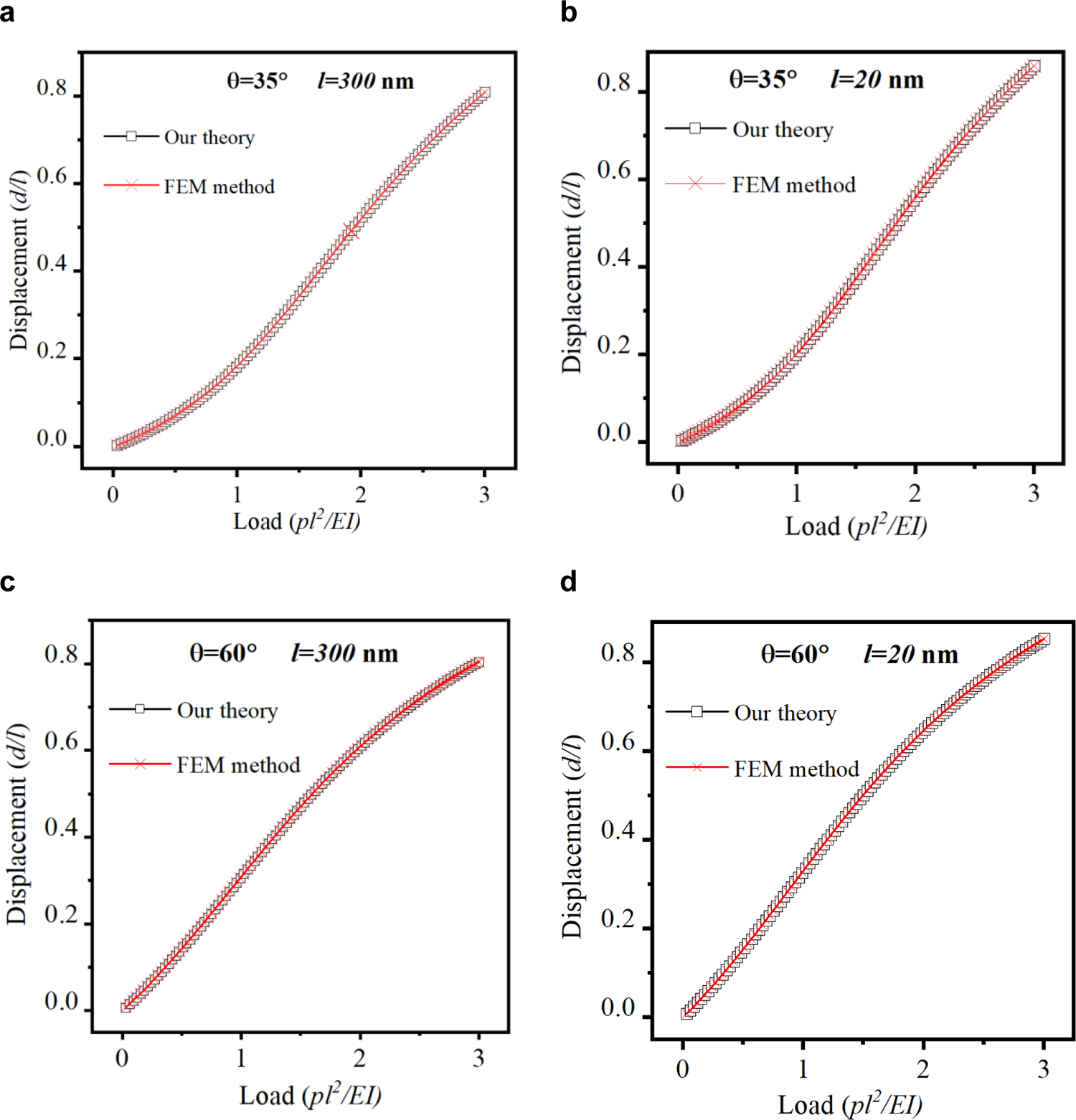
The nonlinear theory is strongly validated by the finite element method (FEM). *_θ_* and *_l_* are inclined angle and length of an actin filament. (**a-d**) Nonlinear deformation behaviours of actin filaments with different lengths and inclined angles.

**Supplementary Fig. 4.**
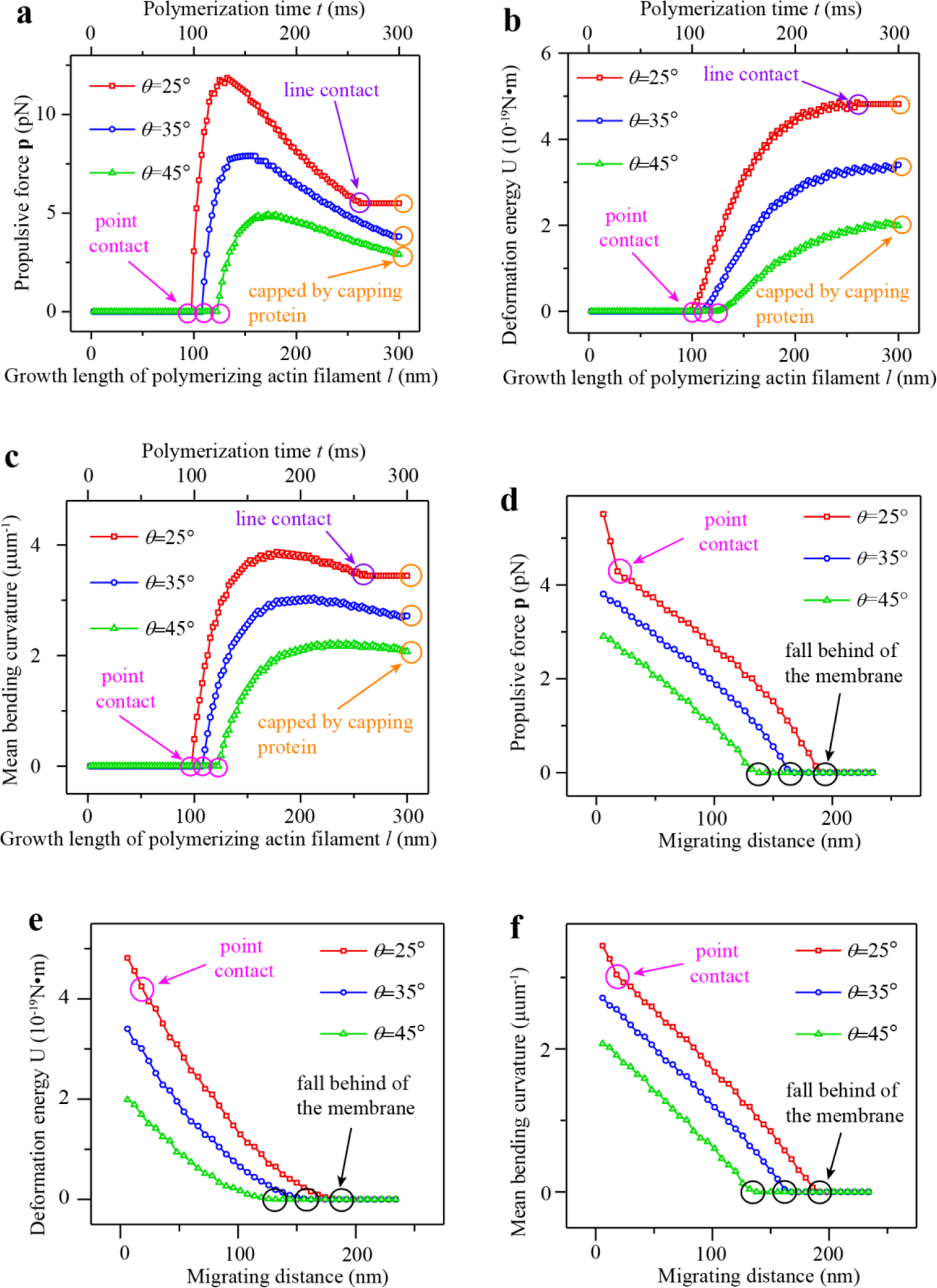
The dynamic interactions between a polymerizing actin filament and the leading-edge membrane during cell migration. (**a-c**) The dynamic evolutions of the propulsive force (**a**), deformation energy (**b**) and mean bending curvature (**c**) when an actin filament is polymerizing toward the leading-edge membrane. (**d-f**) The dynamic evolutions of the propulsive force (**d**), deformation energy (**e**) and mean bending curvature (**f**) when the leading-edge membrane is migrating forward. The pink circle in (**a-c**) is the moment that the barbed end of the polymerizing actin filament begins to attach the leading-edge membrane and the contact between them is a point. The purple circle in (**a-c**) is the moment that the contact between the polymerizing actin filament and the leading-edge membrane changes from a point to a line. The yellow circle in (**a-c**) is the moment that a capping protein caps the barbed end of the polymerizing actin filament and thus the polymerization stops. The pink circle in (**d-f**) is the moment that the contact between the actin filament and the migrating leading-edge membrane changes from a line to a point. The black circle in (**d-f**) is the moment that the actin filament falls behind of the migrating leading-edge membrane and thus they lose contact with each other.

**Supplementary Fig. 5.**
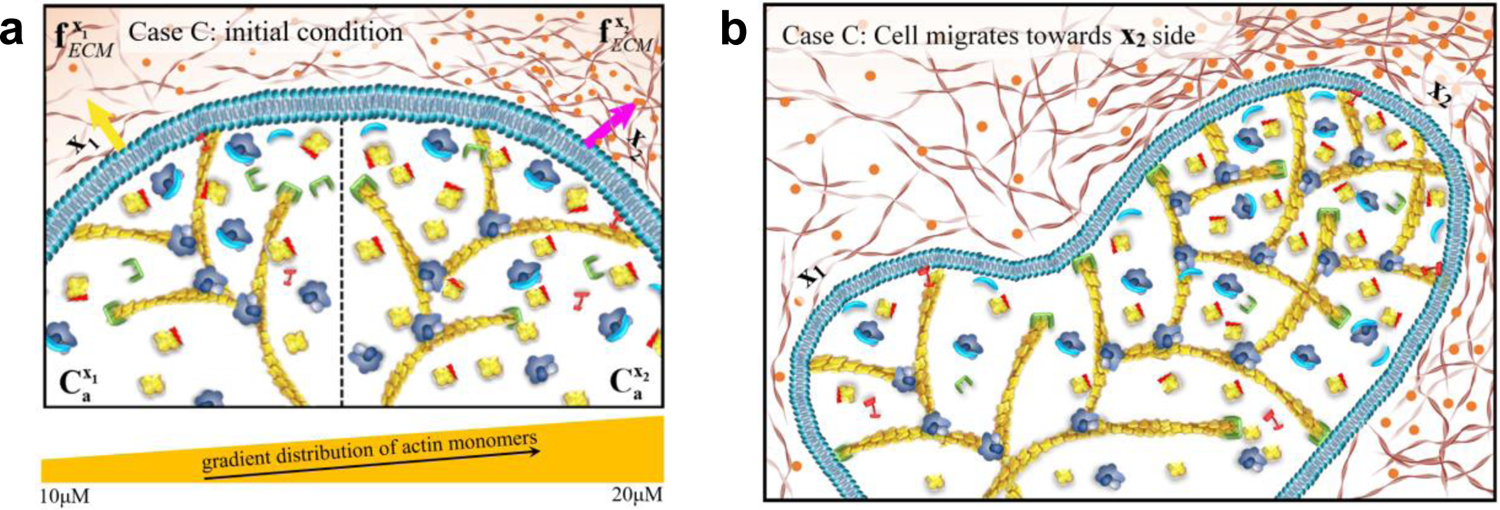
Demonstrations of the initial condition and the simulation results of case C in the directional cell migration section. (a**) Initial condition of case** C. The distribution of actin monomers induced by a chemoattractant has a gradient from the left to the right side. The right-side ECM has a larger resistance. (**b**) Simulation result. It shows that directional cell migration is not solely steered by either the gradients of intracellular proteins induced by chemoattractant or the local stiffness of ECMs. It is a balanced competing consequence between the two.

## Supplementary Information

This supplementary material provides the derivations of the spatiotemporal biophysical theory, and the development of the spatiotemporal multiscale dynamic modelling system.

## Supplementary method

### Derivations of the spatiotemporal “resistance-adaptive propulsion” (RAP) theoretical model

Lamellipodium is one of important ways driving cell migration (Supplementary Fig. 2)(1, 2). Firstly, we investigate the mechanical interactions between a polymerizing actin filament and the curved leading-edge membrane. Near the leading-edge membrane of the lamellipodium, the pointed ends of actin filaments connect with the branched actin network via Arp2/3 complex, which is a relatively rigid protein-protein binding connection(3). The branched actin network has a high stiffness(4, 5) and is fixed by adhesions during cell migration(6). Therefore, the polymerizing actin filaments are assumed to be fixed at their pointed ends. The leading-edge membrane is bent due to the propelling force of the growing branched actin filaments(7). According to the theory of differential geometry, we divide the curved leading-edge membrane into several continuous inclined flat plane segments (Supplementary Fig. 2b). The leading-edge membrane is first assumed to be fixed due to the confinement of the extracellular matrix. When a polymerizing actin filament gets in contact with the membrane, it becomes bent and the interaction force between the bent actin filament and the local plat segment of the membrane is in the direction normal to the flat segment. Though the force interaction between all the bent polymerizing actin filaments and the leading-edge membrane is in three-dimensional space, the interaction between a single polymerizing actin filament and the local flat plane segment of the membrane (i.e., inclined plane) can be described in a two-dimensional deformation plane, which contains the whole bent actin filament (Supplementary Fig. 2c and Fig. 1b in the Main Text).

In our model, the barbed and pointed ends of the polymerizing actin filament are denoted as *s* = _0_ and *s* = _*l*(*t*)_, respectively. Before the polymerizing barbed end reaches the inclined local leading-edge membrane, i.e., *l*_(*t*) cos_ *_β_* < *h*, they have no interaction force (p = _0_) and thus the actin filament is straight. As the actin filament grows in length with the polymerizing time, its barbed end gets in contact with the membrane, i.e., *l*_(*t*) cos_*_θ_* ≥ *h*, the growing filament is compressed and bent due to the constraint from the leading-edge membrane. When the angle *β* (0) of the barbed end is less than *_π_*_/2_, the growing barbed end slides along the membrane in the deformation plane and they have a point contact. The deformed configuration of the actin filament and the interaction force _p_ acting on the leading-edge membrane is shown in Fig. 1b in the Main Text. It is noted that for simplicity, the possible friction between the bent actin filament and the membrane is ignored, thus the force _p_ is always in the normal direction of the local leading-edge membrane. The relation between the bending curvature and the bending moment is expressed as

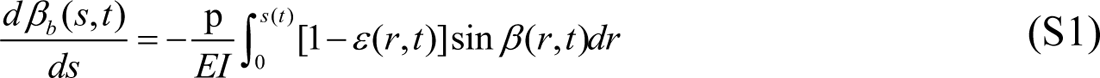

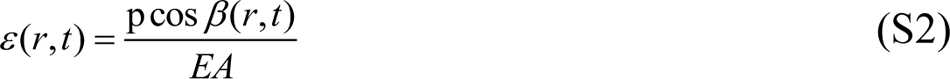

where 0 ≤ *r* ≤ *s*; *E* is Young’s modulus of actin filaments and is taken as 2GPa(8); *A* and *I* are the cross-sectional area and the second moment of the cross-sectional area of the actin filaments, and they can be calculated from the diameter (7 nm) of actin filaments. At the pointed end of the actin filament, i.e., *s* = *l*(*t*), *β_b_* [*l*(*t*),*t*] = *θ*. It is noted that *β_b_* equals *θ* plus the local rotation of the filament resulted from bending, and *β* equals *θ* plus the local rotation of the filament resulted from both bending and transverse shear. Thus, the angle *β_b_* (*s*) along the deformed actin filament as a result of the combined effects of bending, axial compression and polymerization growth can be determined as

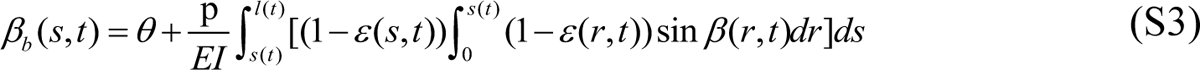

The angle *β_s_* induced by the transverse shear deformation of the polymerizing actin filament is expressed as

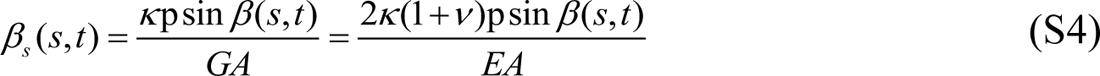

where *v* is the Poisson’s ratio of the actin filament material and is taken to be 0.3(8); and *κ* is the shape factor and is 10/9 for a circular cross-section. Thus, the total angle of the deformed actin filament is

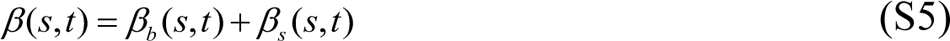

For a polymerization length of the actin filament at time *t*, the nonlinear function *β* (*s*, *t*) can be determined using the iterative method and the following deformation compatibility condition:

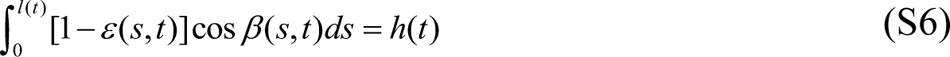

To solve the function *β* (*s*, *t*), we reverse the interacting process by using the local leading-edge membrane (i.e., a plat plane) to compress the actin filament to meet the condition given by equation (S6). More specifically, the actin filament is first assumed to be straight. Then, we divide it into 100000 equal linear segments and apply the load p in the normal direction of the local leading-edge membrane to compress the actin filament at its barbed end, as shown in Fig. 1b in the Main Text, by very small increments of Δp (0.01pN) using an iterative method. The initial axial compressive strain *ε* is chosen as 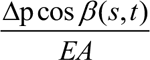 with the initial angle *β* (*s*,*t*) = *θ*. Using the deformation compatibility condition *h*(*t*) in equation (S6), the convergent solution of *β* (*s*, *t*) and the corresponding force p acting on the leading-edge membrane can be obtained by the iterative method. Our nonlinear theory is validated by the finite element method (Supplementary Fig. 5).

The mean curvature *_ξ_* of the geometrically nonlinear deformed actin filament is given as

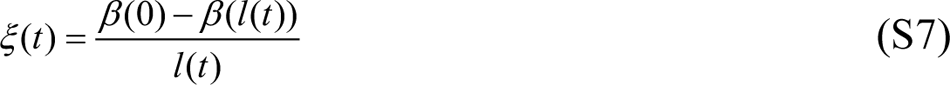

The total elastic strain energy in the actin filament, *U* (*t*), is the summation of the bending energy *U_b_* (*t*), axial compression energy *U_c_* (*t*) and transverse shearing energy *U_s_* (*t*), that is,

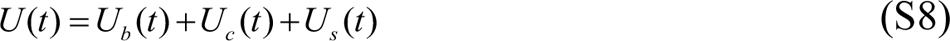

where

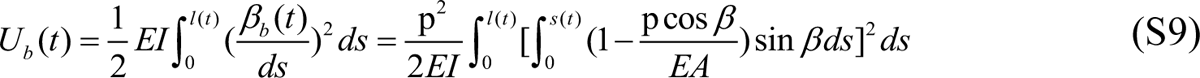

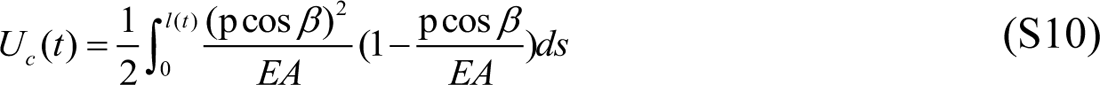

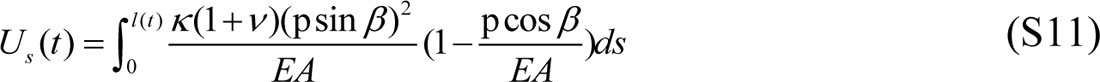

When the angle *β* (0) at the barbed end increases to *π* /2, the barbed end of the actin filament will have a line contact with the local leading-edge membrane (Fig. 1b in the Main Text). Thus, with the continuing polymerization, the actin filament has no further deformation and the interaction force p and the deformation energy *U* keep constant. It is worth mentioning that the interaction force p is in the direction normal to the local leading-edge membrane, rather than in the direction of cell migration. The propulsive force f *_p_* (in the direction of cell migration) acting on the local leading-edge membrane by the actin filament can be expressed by

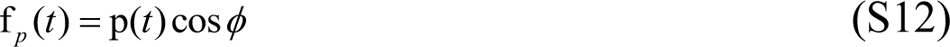

where *ϕ* denotes the angle between the normal direction of the local leading-edge membrane and the direction of cell migration. The dynamic interactions between a polymerizing actin filament and the leading-edge membrane, which is confined by the ECM, are shown in Supplementary Fig. 4a-f. It shows that the polymerizing actin filaments experience large backward nonlinear deformations due to the constraint of the leading-edge membrane and extracellular matrix, which is validated by experimental observations (4, 5, 9), and is also the nature of the actin retrograde flow formation.

Arp2/3 complex has a heterodimer formed by the two subunits, Arp2 and Arp3. The heterodimer bonded with ATP mimics a filamentous actin dimer and thereby creates a template for branching out a daughter actin filament from a mother filament (10, 11). We analyze the conformational deformations of the curved surfaces, both convex and concave sides, of a bending mother filament that potentially interacts with Arp2/3 complex, showing that the convex surface is stretched while the concave surface is compressed (Fig. 1c). Their relative strain *_εr_* relating to the polymerization force-induced bending states can be quantitatively described by *_εr_* _= 2*r*0_*_ξ_* where *_r_*_0_ is the radius of actin filaments (Fig. 1c). Experiments(12) show that *in vitro* long actin filaments (∼10μm) exhibit large bending deformations under thermal fluctuations, and the relative branched density *P*, which is the ratio of the number of branch points with different curvatures to the corresponding number of mother actin filaments, is larger in the convex side than in the concave side. We fit the experimental data of *P* with an inverted sigmoid function *P* = _2 (1+ *e*0.73_*^ξ^*) (Fig. 1d), which is more reasonable than a linear fitting because the relative branched density should be neither minus nor excessively high. The dissociation constant *_Kd_* between Arp2/3 complex and actin filaments is expressed by *K_d_* = [Arp][AF] [ArpAF] where [Arp], [AF], [ArpAF] are the equilibrium concentrations of Arp2/3 complex, actin filaments, and the Arp2/3-actin filament branches, respectively. Since [ArpAF] [AF] = *P*, the dissociation constant becomes *K_d_* = [Arp] *P*. Thus, compared to the straight state of actin filaments, the relative curvature-dependent Arp2/3-actin filament disassociation constant *_K_* (p) *K* ^0^ is

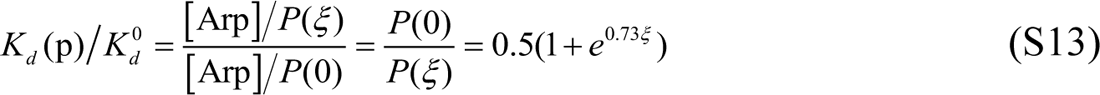

where *K_d_* _(p)_ and *K_d_* ^0^ are the dissociation constants in the bending and straight states, respectively (Fig. 1e). The relative *_K_* (p) *K*_d_^0^ shows that the affinity of Arp2/3 complex binding on the convex surface (negative curvature side) of actin filament is higher than the straight surface, which is higher than the concave surface (positive curvature side). This could be explained by the combination of our mechanical analysis and the recent cryo-electron structure of Arp2/3-actin filament junction(11, 13, 14) that the stretched convex surface of a bending actin filament exposes the groove binding sites for Arp2/3 complex, which facilitate Arp2/3 complex binding, while the compressed concave surface buries some binding sites (Fig. 1c and Supplementary Fig. 1). To better show this force-dependent binding affinity, we also calculate the relative disassociation constant 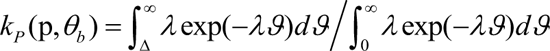 where *K_d_*^1^ (p) and *K_d_* ^2^ (p) are the dissociation constants on the convex and concave surfaces, respectively, demonstrating that the binding affinity on the convex surface is much higher than the concave surface (Fig. 1f). Our theoretical analysis has shown that although *in vivo* actin filaments are relatively short (∼250nm), their polymerization force still can induce them to generate significant backward bending deformations, which could be verified by the actin retrograde flow phenomenon and the recent measurement that the bending curvature can reach up to 10 µm ^-1^ (9). This indicates that the force-dependent actin-Arp2/3 binding affinity occurs *in vivo*. To incorporate that Arp2/3 complex has a higher binding affinity with the convex surface of actin filaments than with straight surface, we introduce a bending curvature-dependent binding factor *_d_ ^arp^* (*_ξ_*^max^) to the model (Eq. S14), which is a segmented function and is defined as the space between two adjacent Arp2/3 complex branches along an actin filament, where *_ξ_* ^max^ is the biggest bending curvature in the deformation history of the actin filament. Since experiments have shown that most of the space *_d_ ^arp^* between two adjacent Arp2/3 branches is 100nm(1, 15), it is appropriate to hypothesize that the relation between *_d_ arp* and the maximum curvature *_ξ_* ^max^ in deformation history follows

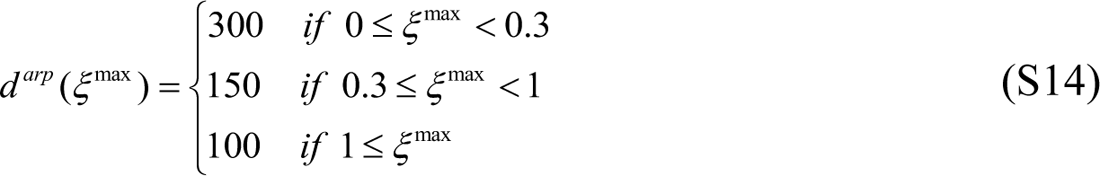

The units of the space *d_arp_* is nm. In the sufficient amount of Arp2/3 complex condition, the number of Arp2/3 complexes binding on an actin filament with a length of *l_i_* can be determined as

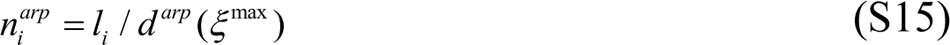

The probabilities that an Arp2/3 complex binding on the convex and concave side of an actin filament are *P*(*ξ*) [*P*(*ξ*) + *P*(− *ξ*)] and *P*(− *ξ*) [*P*(*ξ*) + *P*(− *ξ*)], respectively. Then, Arp2/3 complexes branch out daughter actin filaments from mother actin filaments and thus form the lamellipodial branched actin filament network. During cell migration, the extracellular resistance force is normally in the range of 0.4 nN/μm to 2 nN/μm(16). The leading-edge membrane tension force is about 360 pN/μm(17). The molecular linking attachment force between the barbed ends of actin filaments and the leading-edge membrane, which drag the membrane backward(18–20), is estimated at 20 pN per molecular linker(21–24). The polymerization force of the branched actin network pushes leading-edge membrane to overcome these resistance forces and thus drive the cell to migrate in the ECM.

### Spatiotemporal multi-scale modelling system

Next, we develop a modelling system to spatiotemporally simulate that the three-dimensional lamellipodial self-assembling branched actin filaments drive cells migration forward. In this framework, the interactions between the growing branched actin filaments and the curved leading-edge membrane, which is confined by the varying immediate ECM, are described by the RAP model. The complicated stochastic assembling process and mechano-chemical reactions, such as the highly dynamic actin polymerization, capping protein inhabiting filament growth, Arp2/3 complex nucleation and molecular linker detachment between the filament barbed end and curved leading-edge membrane, are systematically incorporated.

The lamellipodium usually has a thickness of about 200 nanometers and a width of about 20-50 micrometres (25). Thus, we generate a lamellipodial domain of 10, 000 nm × 1, 000 nm × 200 nm and use periodic boundaries in the width direction. During cell migration, the propulsive force of the actin filaments bends leading-edge membrane(7). By referring to the differential geometry theory, we divide the smoothly curved leading-edge membrane into *_N_* inclined flat plane segments **P** = [*p*_1_, *p*_2_, *p*_3_] as shown in Supplementary Fig. 2. The *k*th membrane plane has a point with coordinates of **x** *^p^_k_* and a normal vector **x** *^pn^_k1_*. Let **x** *^p^* and **x** *^pn^* be the concatenations [**x** *^pT^_1_*, **x** *^pT^_2_*, …… **x** *^pT^_N_*] and [**x** *^pn T^_1_*, **x** *^pn T^_2_*, …… **x** *^pn T^_N_*]*^T^*, respectively. During lamellipodium driving cell migration, the leading-edge protrude forward in the extracellular matrix. Thus, the equations **_M_**_(**P**, *t*)_ of these inclined flat plane segments **P** in three-dimensional space can be written as

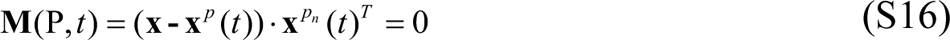

In our simulations, the leading-edge membrane is divided into six inclined planes. Then, we generate a certain concentrations of actin monomers C*_a_*, Arp2/3 complex C*_arp_*, capping proteins and ATP in the lamellipodial space. The distribution of actin monomers is defined by a gradient extracellular chemoattractant (Supplementary Fig. 5). These actin monomers stochastically nucleate into a certain number *n* of short actin filaments, and the pointed end coordinates **x** *^fp^* of the *i*th actin filament are randomly and independently generated. For each an actin filament growing in the lamellipodial sheet-like space, a local spherical coordinate system is created by regarding its pointed end as the origin. This local spherical coordinate system is used to determine the orientation and the barbed end coordinates (*r ^fb^, φ ^fb^, θ ^fb^*) of each actin filament confined in the lamellipodium. Published experimental results showed that the orientation of actin filaments relative to the cell migration direction is generally around ±35° (26). Thus, the orientation angles of the initial actin filaments are randomly produced by a normal distribution and a uniform distribution defined as

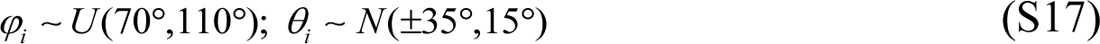

The behaviour of actin is coupled to the nucleotide bound to its active site, and ATP hydrolysis is activated during polymerization(27). Thus, the polymerization process of actin filaments is simulated by adding individual actin monomers bonded by an ATP to the barbed ends. The growth rate *V_fil_* of the polymerizing free barbed end of an actin filament is expressed as

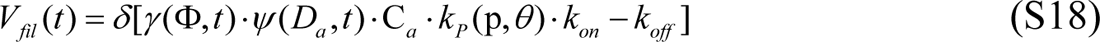

where *_δ_* is the radius of an actin monomer; C*_a_* is the local concentration of actin monomers in the cell; *γ* (Φ) is the consuming factor of actin monomers, introducing the relation that polymerizing rate is proportional to the ratio of the concentration of actin monomers to the density of polymerizing filaments _Φ_ (28); *_t_cap* and *_t_cap* are the nucleation and capping time of the actin filament, respectively. *k_P_* (p,*θ*) is an interacting force-induced probability density distribution in the rate of filament polymerization due to size-dependent insertion of monomers 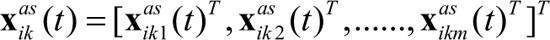 (29, 30). p is the interaction force that the polymerizing actin filament acts on the leading-edge membrane; *θ_b_* is the angle of the filament barbed end relative to the normal vector of the local membrane; *_ϑ_* is a function of p and *θ_b_* and Δ is the size of sufficient gap to permit intercalation of monomers (29, 30). *k_on_* and *k_off_* are the polymerization and depolymerization rate constants, respectively. p and *θ_b_* are calculated based on the RAP theory. *ψ* is a scaled diffusion coefficient of actin monomers and is to introduce the effect of actin filament density on the actin diffusion flux to the polymerizing barbed ends based on the Fick’s first law of diffusion. *D_a_* is the diffusion coefficient of actin monomers for supplying the polymerization of the leading-edge actin filaments, and is described as

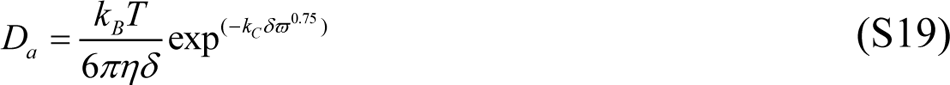

in which *k_B_* is Boltzmann’s constant, *T* is temperature, *η* is the viscosity of water at *T*, *ϖ* is the volume fraction of branched actin filament near the leading edge, and *k_C_* is a regressed constant(31). Let **_x_** *^fp^* be the concatenation [**x** *^fpT^*, **x** *^fpT^*,, **x** *^fpT^*]*^T^*. Thus, before the barbed ends attach the leading-edge membrane, their coordinates **x** *^fb^* (*t*) at time *t* are specified by

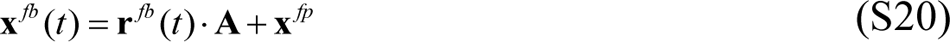

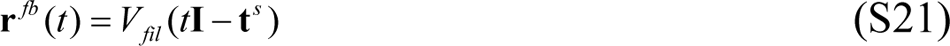

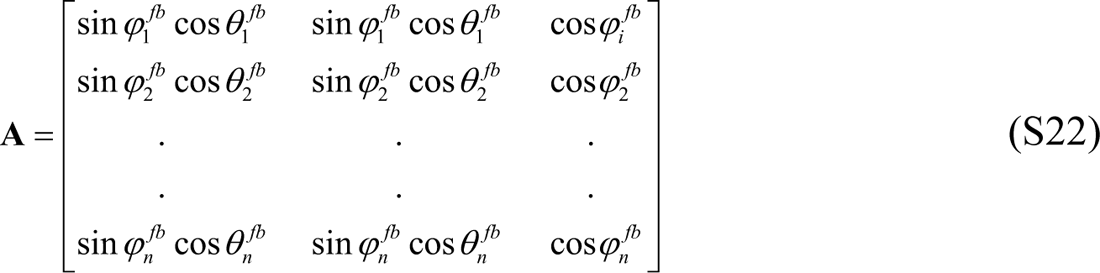

where **I** denotes the identity matrix and **t***^nuc^* = *diag*(*t^nuc^, t^nuc^*,,*t^nuc^*). is the time when the *i*th actin filament nucleates and begins to polymerize. In lamellipodia, actin filaments can normally grow to a length of about 200 nm(1, 32). Thus, we assume that when an actin filament grows to its maximal length, which is generated by the normal distributions *N*(200,50), capping proteins will cap its barbed end and the polymerization will stop. Thus, the final lengths of actin filaments can also be expressed by

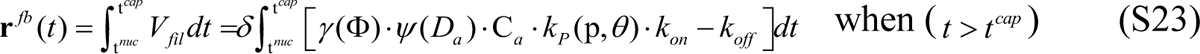

where **t***^cap^* = *diag*(*t^cap^, t^cap^*,……, *t^cap^*) with *t^cap^* being the time that a capping protein binds on the barbed end of the *i*th actin filament. The final coordinate *^fb^* of its barbed end in the lamellipodial sheet-like space should be within the domain given by

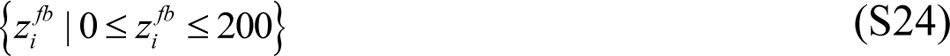

Once their barbed ends reach the leading-edge membrane, the spatiotemporal mechanical interaction between the polymerizing actin filaments and the bent protruding leading-edge membrane can be described by our RAP theoretical model. The whole process of lamellipodia protrusion is in a highly dynamic state. N-WASP proteins form molecular linkers between the barbed ends of actin filaments and the leading-edge membrane. Meanwhile, Arp2/3 complexes are activated by Wiskott-Aldrich syndrome proteins (WASPs). Then, they bind on actin filaments and branch out new actin filaments. The probabilities of the Arp2/3 complex binding on the convex and concave sides of an actin filament are *P*(|*ξ*|) [*P*(|*ξ*|) + *P*(− |*ξ*|)] and *P*(− |*ξ*|) [*P*(|*ξ*|) + *P*(− |*ξ*|)], respectively. To identify the binding coordinates of the Arp2/3 complex along actin filaments during this dynamic process, we create a series of local Cartesian coordinate systems (*x*′, *y*′, *z*′) by regarding the pointed end of each actin filament as the origin *_O_*_′_. Plane. *O*′*x*′*y*′. is the deformation plane of the bent actin filament and axis *_O_*′*_x_*′ is the normal vector of the inclined leading-edge membrane flat plane segment. For the *j*th Arp2/3 complex on the *i*th actin filament, which pushes against the inclined leading-edge membrane plane segment *_k_*, the deflection vector Δ**x**′*_ikj_* (*t*) of the binding point of the Arp2/3 complex in the local coordinate system *O_i_*′*_k_ x_i_*′*_k_ y_i_*′*_k_ z_i_*′*_k_* at time *t* can be calculated with the RAP theoretical model. If there are *m* Arp2/3 complexes binding on the *i*th actin filament, let. Then, with Jacobian determinant **J**, the dynamic coordinates of the start point of Arp2/3 complex **x***^as^* (*t*) = [**x***^as^* (*t*)*^T^*, **x***^as^* (*t*)*^T^*,……, **x***^as^* (*t*)*^T^*]*^T^* in the global Cartesian coordinate system at time *t* can be obtained as

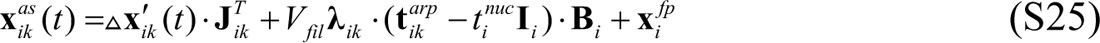

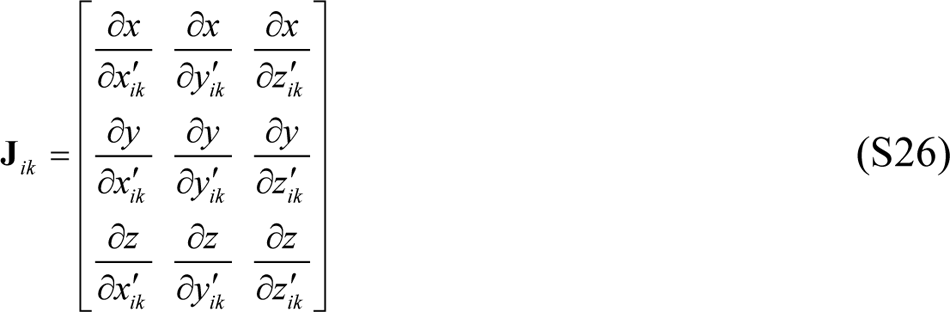

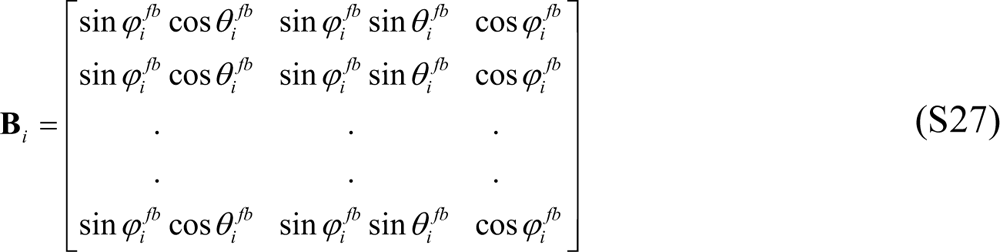

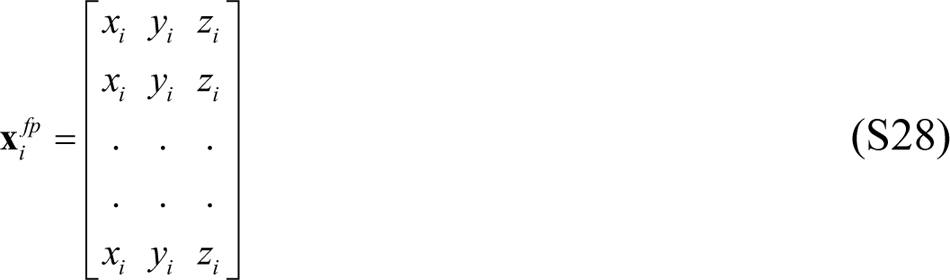

where and denotes the time that the *j*th Arp2/3 complex attaches the *i*th actin filament due to the mechanical interaction with the local leading-edge membrane plane segment is randomly generated in the range of (0, 1). Matrices **B***_i_* and **I***_i_* all have *m* rows. The Jacobian determinant is calculated from the transformation relationship between the local (*x_i_*′*_k_ y_i_*′*_k_ z_i_*′*_k_*) and global (*xyz*) Cartesian coordinate systems.

The length of Arp2/3 complex *r^arp^* is normally about 10 nm (33). The subunits Arp2 and Arp3 bind ATP via nucleotide-binding domains, and then assemble branched actin network through hydrolysis of ATP(34). During this process, Arp2/3 complex binds on the convex side of a bent actin filament (12) with an angle of ∼70°(3). Thus, the possible branching position of Arp2/3 complex is constrained on a half conical surface around the actin filament. The azimuthal angle of Arp2/3 complex in its local spherical system is also generated by a uniform distribution *U* (70°,110°). Because the forthcoming daughter actin filament will grow out from the Arp2/3 complex, angle *φ ^arp^* also should allow the daughter filament to polymerize to a length *r ^df^* specified by a normal distribution *N*(200,50). Therefore, the azimuthal angle and the end point coordinates **x***^ae^* (*t*), i.e., the vertical vector [*x^ae^* (*t*), *y^ae^* (*t*), *z^ae^* (*t*)], of the *j*th Arp2/3 complex on the *i*th actin filament interacting with the local leading-edge membrane segment *k* at time *t* should satisfy

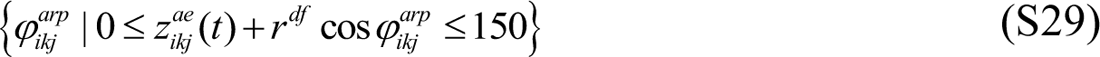

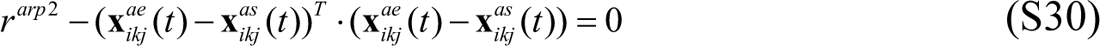

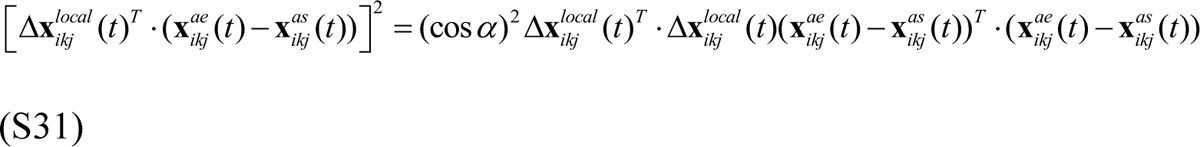

where Δ**x***^local^_ikj_* (*t*) is the local tangent vector of the bent actin filament segment where Arp2/3 complex binds. This vector updates with time because of actin filament polymerization, leading-edge membrane protrusion and their dynamic interaction, and can be calculated with the RAP model. *α* denotes the angle between the mother filament and the Arp2/3 complex, and is generated by a Gaussian distribution of *N*(70°, 2°). Thus, the end point coordinates **x***^ae^_ikj_* (*t*) of Arp2/3 complex can be specified and updated by using the equations (S29-31). After the binding of Arp2/3 complex, daughter actin filaments will nucleate there. The concatenation matrix of the pointed ends of daughter actin filaments 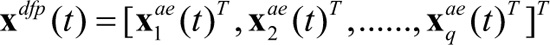 and *q* is the total number of Arp2/3 complex. Then, the daughter filaments polymerize by adding actin monomers onto their barbed ends in the branching direction of the Arp2/3 complex. The positions of the daughter filaments are also in a dynamic state with Arp2/3 complex, and can be determined by the deflection equations of their mother filaments calculated from the RAP model. Like the mother actin filaments, the polymerizing daughter filaments will also mechanically interact with the leading-edge membrane and be bent. Their spatiotemporal positions are updated by the RAP model. New activated Arp2/3 complex by

WASPs will also bind on these daughter filaments and branch out next generation daughter actin filaments. With the increase in the number of actin filaments pushing against the leading-edge membrane, the leading-edge membrane will protrude forward. Consequently, these bent actin filaments will straighten, either partially or totally. Meanwhile, the assembling and mechano-chemical reactions between these different proteins are constantly taking place, remodeling and organizing new lamellipodial cytoskeleton to drive cell migrations in ECMs. References

---

**Supplementary Table 1.**
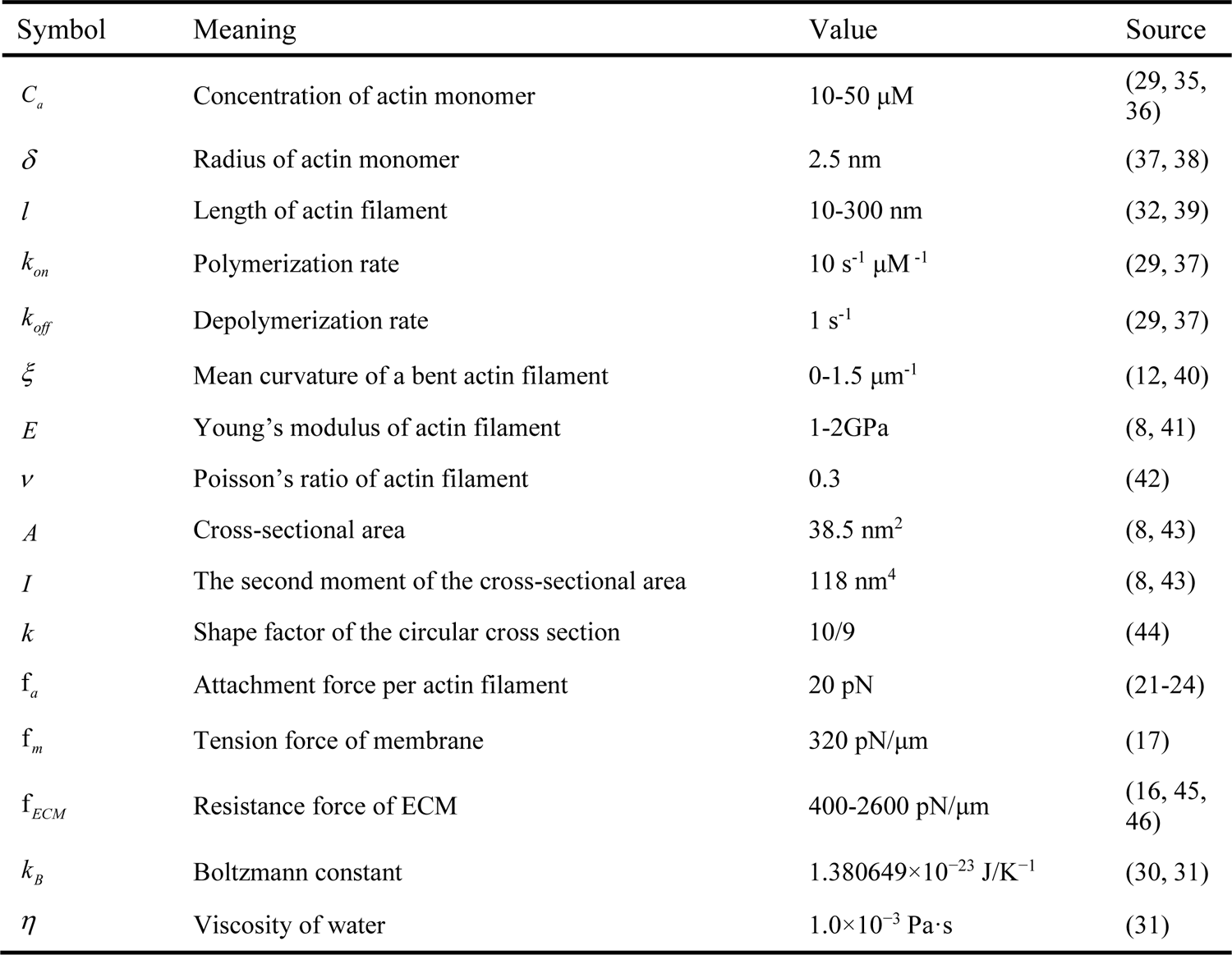
Parameter values

| Symbol | Meaning | Value | Source |
| --- | --- | --- | --- |
| $C_a$ | Concentration of actin monomer | 10-50 $\mu\text{M}$ | (29, 35, 36) |
| $\delta$ | Radius of actin monomer | 2.5 nm | (37, 38) |
| $l$ | Length of actin filament | 10-300 nm | (32, 39) |
| $k_{on}$ | Polymerization rate | 10 $\text{s}^{-1} \mu\text{M}^{-1}$ | (29, 37) |
| $k_{off}$ | Depolymerization rate | 1 $\text{s}^{-1}$ | (29, 37) |
| $\xi$ | Mean curvature of a bent actin filament | 0-1.5 $\mu\text{m}^{-1}$ | (12, 40) |
| $E$ | Young's modulus of actin filament | 1-2GPa | (8, 41) |
| $\nu$ | Poisson's ratio of actin filament | 0.3 | (42) |
| $A$ | Cross-sectional area | 38.5 $\text{nm}^2$ | (8, 43) |
| $I$ | The second moment of the cross-sectional area | 118 $\text{nm}^4$ | (8, 43) |
| $k$ | Shape factor of the circular cross section | 10/9 | (44) |
| $f_a$ | Attachment force per actin filament | 20 pN | (21-24) |
| $f_m$ | Tension force of membrane | 320 pN/$\mu\text{m}$ | (17) |
| $f_{ECM}$ | Resistance force of ECM | 400-2600 pN/$\mu\text{m}$ | (16, 45, 46) |
| $k_B$ | Boltzmann constant | 1.380649$\times 10^{-23}$ J/K$^{-1}$ | (30, 31) |
| $\eta$ | Viscosity of water | 1.0$\times 10^{-3}$ Pa s | (31) |

**Supplementary Table 2.**
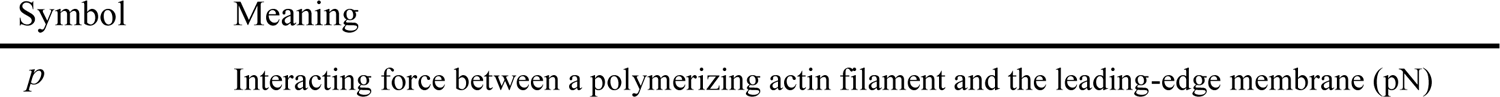
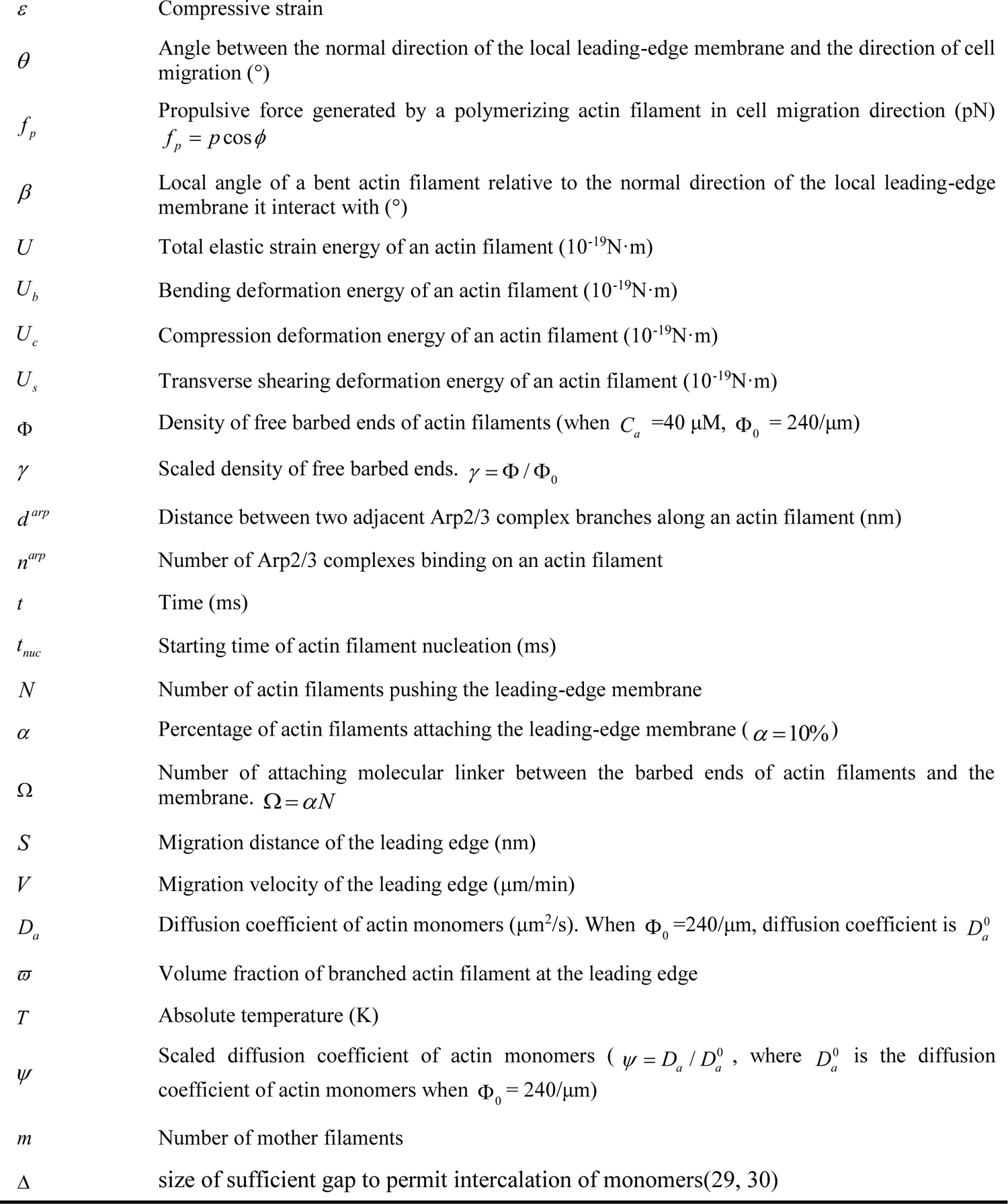
Other notations

